# Rare twin cysteine residues in the HIV-1 envelope variable region 1 link to neutralization escape and breadth development

**DOI:** 10.1101/2024.09.05.611179

**Authors:** Maria C. Hesselman, Marius Zeeb, Peter Rusert, Chloé Pasin, Jennifer Mamrosh, Samuel Kariuki, Michèle Sickmann, Masako M. Kaufmann, Daniel Schmidt, Nikolas Friedrich, Karin J. Metzner, Audrey Rindler, Herbert Kuster, Craig Adams, Ruwayhida Thebus, Michael Huber, Sabine Yerly, Karoline Leuzinger, Matthieu Perreau, Roger Koller, Günter Dollenmaier, Simona Frigerio, Dylan H. Westfall, Wenjie Deng, Allan DeCamp, Michal Juraska, Srilatha Edupuganti, Nyaradzo Mgodi, Hugh Murrell, Nigel Garrett, Kshitij Wagh, James I. Mullins, Carolyn Williamson, Penny L. Moore, Huldrych F. Günthard, Roger D. Kouyos, Alexandra Trkola

## Abstract

The identification of HIV-1 Envelope glycoprotein (Env) traits associated with development of neutralization cross-reactivity in natural infection is critical for vaccine design. Here we describe the presence of additional Cysteine (Cys) residues in V1 that are enriched among people with elite neutralization breadth. Using >65,000 V1 sequences from the CATNAP database, the AMP trials and three large longitudinal HIV infection cohorts, the SHCS, ZPHI and CAPRISA studies, we show that Env variants with extra V1 Cys are present at low levels throughout infection and fluctuate in frequency over time within participants. We demonstrate an independent association of extra V1 Cys with elite plasma neutralization, and a strong preference for two versus one extra Cys, suggesting certain Envs introduce an additional disulfide bond for stabilization. We observed high levels of neutralization resistance among Envs from 34 bNAb donors, of which 17.6% had elongated V1 regions with extra Cys. We show that extra V1 Cys moderately increase neutralization resistance in an Env from a V2- Apex bNAb-inducer. Modulation of the accessibility of bNAb epitopes on this Env by extra V1 Cys enhanced epitope shielding of several regions, but increased V2 exposure. This suggests that escape from autologous neutralizing activity drove insertion of the extra V1 Cys, creating a modified antigen that may have favored V2 bNAb induction in this donor. Overall, we identify a rare motif of twin Cys in V1 that confers increased neutralization resistance and Env stabilization, is associated with bNAb induction, and may hold potential for incorporation into future HIV bNAb immunogens.

## Introduction

Genetic traits of the HIV-1 envelope (Env) that link to bNAb development and escape in untreated HIV-1 infection are key to HIV-1 vaccine immunogen development. A known regulator of neutralization resistance is the Env region encompassing variable loops 1 and 2 (V1V2), which affects the neutralization properties of multiple epitopes including those of bNAbs^1–10^. Spanning residues 126 to 196 in the gp120 portion of Env (HXB2 numbering), the V1V2 domain is positioned at the apex of the Env trimer^11–13^. There, it forms a top layer on the trimer that shields key neutralization sensitive sites, the variable loop 3 (V3) and the co-receptor binding site.

The high degree of sequence and length heterogeneity observed in the V1V2 is indicative of its remarkable adaptability^1–10^. Longer and more glycosylated V1V2 can decrease susceptibility to many bNAbs^1,4^. Furthermore, V1V2 length dynamics during infection align with the presence or absence of neutralizing antibodies (nAbs)^1,3^. During sexual transmission of HIV-1, a robust genetic bottleneck allows only a few transmitted variants to establish the infection^14–17^, and in some transmission cohorts, shorter and less glycosylated V1V2 regions appeared to be favored during transmission and early infection^3,18^. The onset of humoral immunity and nAb development sees an elongation of the variable loops which subsides with waning humoral immunity in late infection stages^3^. While longer V1V2 regions may provide an advantage in the presence of nAbs, the shortening of the loops in environments where nAbs are absent highlights the potential detrimental effect of V1V2 elongation on viral fitness.

We previously conducted the Swiss 4.5K screen^19,20^, a population-level investigation of bNAb activity in 4484 people with HIV (PWH) participating in the Swiss HIV Cohort Study (SHCS)^21,22^ and the Zurich Primary Infection Study (ZPHI)^23^. The Swiss 4.5K screen led to the definition of key parameters of bNAb induction and identified 239 bNAb-inducers^19,20^. This screen and a follow-up study investigating 303 transmission pairs revealed a clear effect of virus genetics on the antibody response^24^. Here, building on these findings, we conducted a search for Env genetic and phenotypic features that are enriched among bNAb-inducers.

## Results

### High neutralization resistance among envelope glycoproteins from bNAb-inducers

To gain insight into global Env features associated with bNAb induction and escape, we leveraged a collection of bNAb-inducer samples and data identified in the Swiss 4.5K screen screen^19,20^. We created a panel of *envs* cloned from chronic stage plasma virus (≥2 years post estimated date of infection, EDI) of 34 bNAb inducers with diverse neutralizing specificities and high breadth and potency (Figure S1A, Table S1). Infection competent Envs, one per participant, were assembled into a bNAb-inducer Env pseudovirus (PSV) panel (N=34) spanning several subtypes (Figure 1A, Table S2). With the exception of two Envs from a previously identified bNAb transmission pair (S30402 and S30349)^24^, the 34 bNAb-inducer *envs* were not related (Figure 1A).

**Figure 1:**
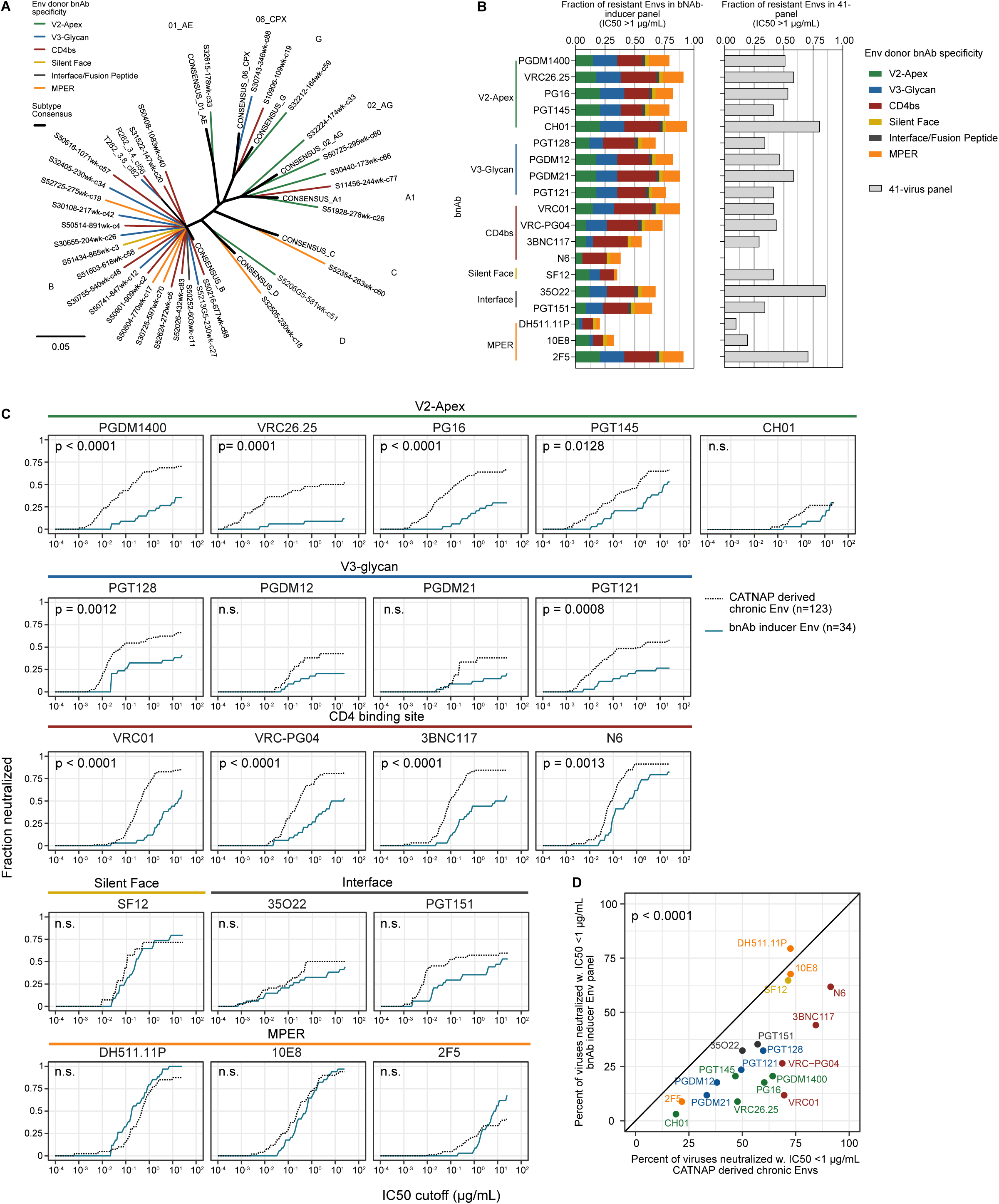
bNAb-inducer Envs have a generalized neutralization resistance. (A) Phylogenetic tree of *env* nucleotide sequences of bNAb-inducer Env panel (N=34 Envs) (Table S2). Branch color indicates the predicted plasma bNAb specificity of the donor. Env subtype consensus sequences (2005 version) from the Los Alamos HIV Sequence Database are shown as thicker black lines. (B) Resistance of the bNAb-inducer Env panel (N=34) to known bNAbs (left) and multiclade 41-panel (right). Resistance to each bNAb is quantified as the fraction of Envs with a half-maximal inhibitory concentration (IC50) >1 µg/mL. Colors indicate the predicted plasma bNAb specificity of the donor. (C) Cumulative frequency distribution (breadth-potency curves) of the IC50 values of 19 bNAbs against the bNAb-inducer Env panel (blue) and CATNAP-derived chronic Envs (black dotted line) (123 Envs sampled in chronic infection). P-values are derived from Wilcoxon test by comparing median IC50 of each bNAb against the bNAb-inducer and CATNAP-Envs (Figures S1D and E). (D) Comparison of the percentage of highly sensitive Envs (IC50 < 1 µg/mL) in the CATNAP chronic Env (x-axis) and bNAb-inducer Env panels (y-axis) for each of the 19 bNAbs tested. P-value calculated with t-test comparing mean breadth of all antibodies between the two panels.

We measured sensitivity of the bNAb-inducer PSV panel to reference bNAbs (N=19) covering all major bNAb epitopes including some of the most potent and broad bNAbs identified (Table S3). The sensitivity of the bNAb-inducer PSV panel was significantly lower to 15/19 reference bNAbs (Wilcoxon p<0.05) when compared to a previously described^19^ multi-subtype panel of Envs (N=41; Table S4) from acute and chronic infection that we routinely use for bNAb breadth analysis (Figure S1B and S1C). We expected to observe epitope-specific bNAb resistance, corresponding to *in vivo* escape from the bNAb specificity in the donor. Instead, we observed a remarkable generalized resistance of bNAb-inducer Envs to neutralization, extending beyond potency limitations of the bNAbs, with the majority of bNAb-virus combinations having 50% inhibitory concentrations (IC50) >1 µg/mL (Figure 1B, Table S2). Notably, four bNAbs in clinical development, VRC01, 3BNC117, PGT121 and PGDM1400, all showed poor neutralization breadth against bNAb-inducer Envs. Only the CD4 binding site (CD4bs) bNAb N6, the membrane proximal region (MPER) bNAbs DH511.11P and 10E8 and the silent face (SF) bNAb SF12, retained high breadth against the resistant bNAb-inducer Envs, neutralizing more than 50% of Envs with IC50s ≤1 µg/mL which was not the case for the 41-virus panel (Figure 1B). Resistance to V3-glycan and V2-Apex bNAbs was particularly high despite similar frequencies of key glycosylation sites for V3-glycan (N332/N334) and V2-Apex (N156/N160) bNAbs among bNAb-inducer Envs and the 41-virus panel (Tables S2 and S4).

Because the multi-subtype 41-Env panel included several Envs from acute infection (Table S4) whereas Envs in the bNAb-inducer PSV panel were all derived from chronic infection, we compared the bNAb-inducer Envs to a separate set of 123 chronic infection Envs, using data available in the CATNAP^25^ database (Table S5). This analysis confirmed that bNAb-inducer Envs were more neutralization resistant than average Envs from chronic infection. Setting IC50 < 1 µg/mL as a potency threshold, bNAbs showed markedly lower breadth against bNAb-inducer Envs compared to CATNAP-chronic Envs (Figure 1D, t-test p<0.0001). Ten out of 19 bNAbs tested were significantly less potent against the bNAb-inducer Env panel than CATNAP-chronic Envs (Wilcoxon p-values <0.0001 to 0.03) (Figures 1C, S1D-E) and several more bNAbs (CH01, PGDM12, PGDM21, PGT151) with overall lower breadth showed the same trend without reaching significance. Only the MPER bNAbs, the SF bNAb SF12 and the interface bNAb 35O22 had similar potency against bNAb-inducer Envs (Figures 1C, S1D-E). Since the bNAb-inducer panel was enriched for subtype B viruses (22/34; Table S2), we conducted a subtype B-restricted comparison which confirmed its higher resistance compared to CATNAP-Envs of subtype B (Figure S1E).

Collectively, we conclude that bNAb-inducer Envs have a marked generalized neutralization resistance against various bNAbs that is not subtype specific and extends beyond bNAb-specific escape.

### Distinct features of the V1 hypervariable loop region among bNAb-inducer Envs

Env variable loop length and glycosylation can act as a regulator of neutralization sensitivity^1–10^. Comparing variable loop lengths (V1-V5) between the bNAb-inducer and the CATNAP- chronic Env panels, we observed significantly longer V1, V2 and V5 loops in bNAb-inducer Envs (Figure 2A). The largest difference was in V1 (mean length 35 vs 29 residues, t-test p < 0.0001) (Figure 2A). The differences in V2 and V5 length were less pronounced, though statistically significant, whereas V3 and V4 lengths were not significantly different between the bNAb-inducer and CATNAP-chronic Env panels (Figure 2A). Furthermore, bNAb-inducer Envs had on average more glycosylation sites in V1, V2 and V3 loops than CATNAP-chronic Envs and V4, V5 loops had fewer glycosylation sites (t-test p<0.05) (Figure 2B).

**Figure 2:**
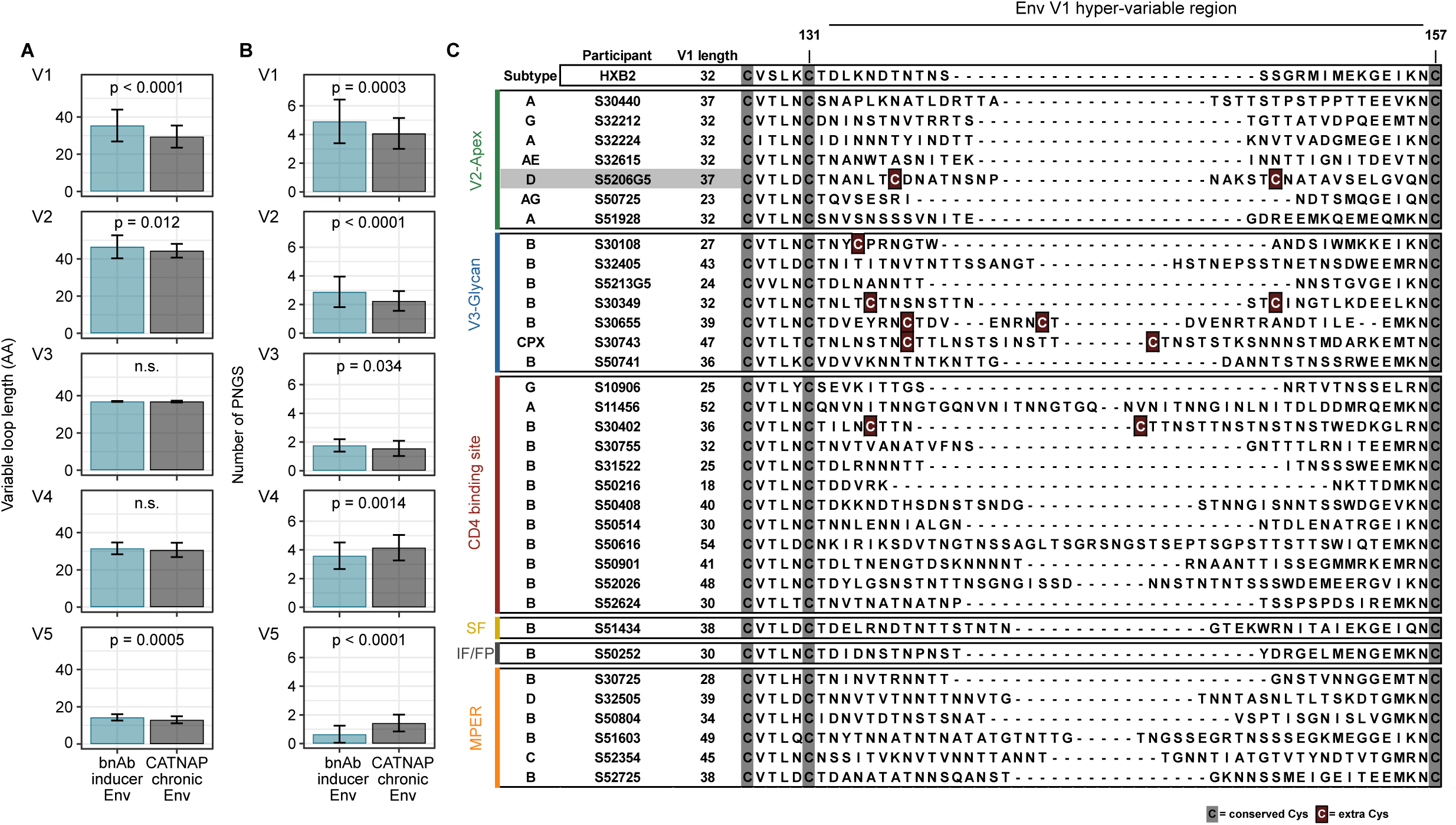
bNAb-inducer Envs have longer V1 loops with non-canonical Cysteine residues. (A) Comparison of variable loop length (V1-V5) of bNAb-inducer Envs (n=34) CATNAP- derived chronic Envs (n=123). P-values calculated with t-test on the mean length in each group. (B) Comparison of the number of predicted N-linked glycosylation sites (PNGS) in V1-V5 between bNAb-inducer Envs (n=34) CATNAP-derived chronic Envs (n=123). P-values calculated with t-test on the mean number of PNGS in each group. (C) Multiple sequence alignment of the V1 hyper-variable region (HXB2 residues 126-157) of the bNAb-inducer Env panel. Gray and burgundy boxes indicate conserved and extra V1 Cys, respectively. The predicted specificity of the bNAb inducer plasmas from which the Envs were isolated is indicated on the left.

In the context of the closed Env trimer conformation, the V1V2 region forms a Greek key motif held together by disulfide bonds between two pairs of highly conserved cysteine (Cys) residues^26,27^, one in V1 between residues 131 and 157 and a second in V2 between residues 126 and 196^28^. Analysis of the V1 regions of the bNAb-inducer Env panel showed that 6/34 bNAb-inducer Envs (17.6%) had non-canonical Cys residues in their V1 loops in addition to the conserved Cys at 131 and 157 (Figure 2C). Insertions of two Cys residues were preferred over single Cys insertions, with five of the bNAb-inducer Envs bearing a twin Cys insertion, whereas only one had a single Cys insertion. The five Envs with a twin Cys insertion had especially long V1 regions averaging 38 residues, compared to an average of 29 residue long V1 regions among CATNAP-chronic Envs. Extra V1 Cys were found in Envs of different subtypes (B, CPX, D) and from donors with different predicted bNAb neutralization specificities (V3-glycan, CD4bs, V2-Apex) with 4/6 Envs stemming from V3-glycan bNAb activity predicted donors (Figure 2C).

The preferential addition of twin Cys suggests that an additional disulfide bond may form in these Envs, which may contribute to the conformational stability of the region. The presence of twin Cys residues in elongated V1 loops has been reported thus far only in few donors, notably including seven cases of bNAb-inducers^6,29–31^. Taken together, the observation of V1 Cys insertions in bNAb-inducer Envs by us and others suggests that V1 Cys insertions may be associated with bNAb evolution.

### Cys insertions in V1 promote low-level neutralization resistance

We hypothesized that stabilization of the V1 domain through an additional disulfide bond may contribute to the high degree of generalized neutralization resistance in some bNAb-inducer Envs. To investigate this, we followed plasma neutralization and *env* evolution longitudinally in donor S5206G5, an elite neutralizer with a predicted V2 apex bNAb specificity (Table S6)^19^. The S5206G5 Env (S5206G5-581wk-c51) included in the bNAb-inducer Env panel combines the features we observed: a long V1 region (37 residues) including a twin V1 Cys insertion and high resistance to a wide range of bNAb types (Figure 2C, Table S2). S5206G5 plasma neutralization breadth against a 19-virus multiclade panel, peaked at 90% in week 610 post estimated date of infection (EDI) and reached a maximum geometric mean 50% neutralization titer (NT50) of 1:6’275 at week 643 (Figure S2A). Sequence analysis of *env* cloned from plasma viruses at 12 time points revealed that variants with no extra V1 Cys (C^0^), one extra V1 Cys (C^1^) and two extra V1 Cys (C^2^) co-existed at week 42, the earliest available sample (Figure 3A, Figures S2B-C). Afterwards, the C^2^ motif became predominant and the V1 region longer, suggesting that this motif provided a benefit for the virus (Figure 3A).

**Figure 3:**
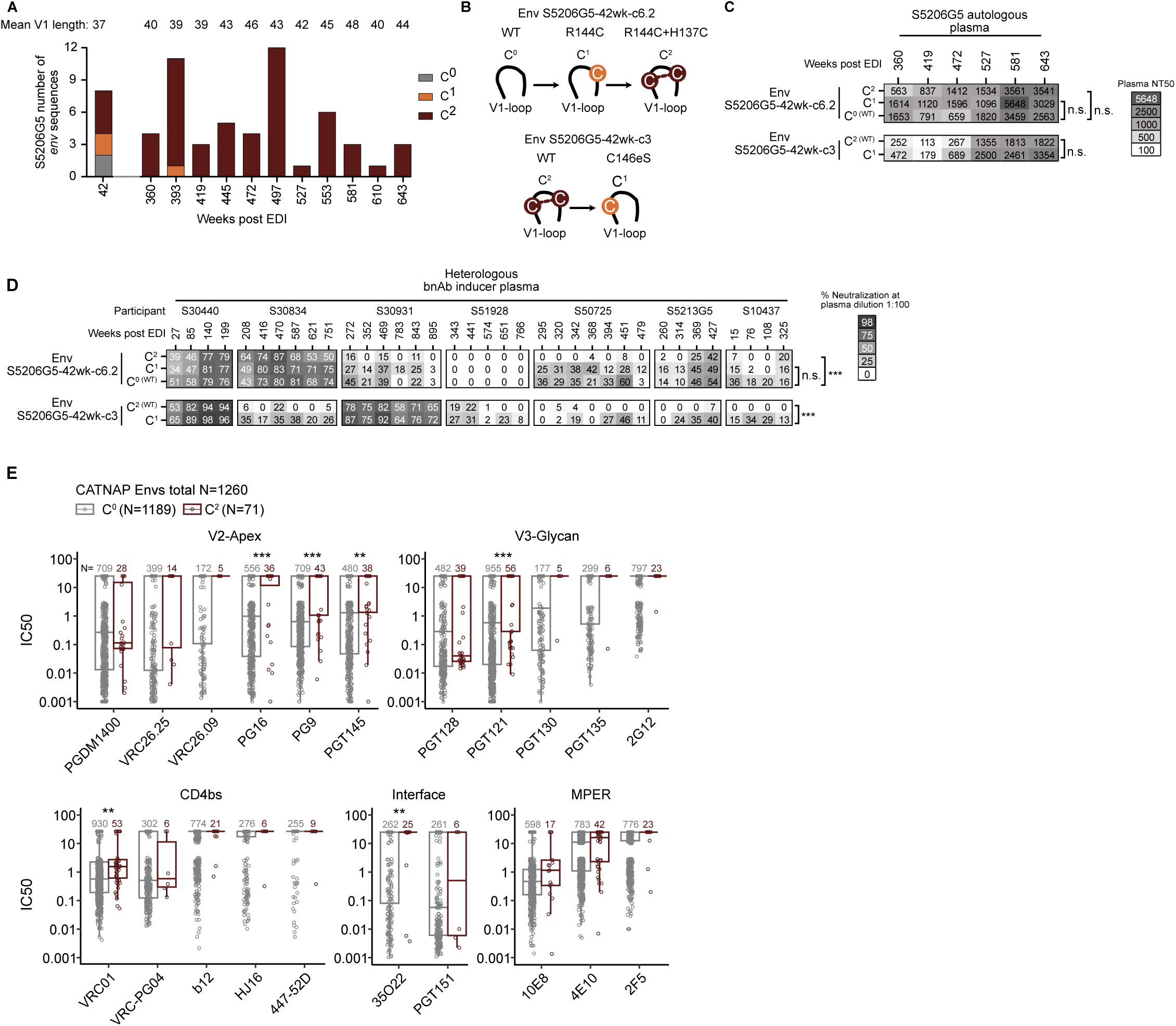
Effects of V1 Cys insertion in bNAb-inducer S5206G5 Env on plasma neutralizing activity. (A) Number of S5206G5 Env variants with extra V1 Cys per sample time point (42-643 weeks post EDI). Mean V1 length of each time point indicated above. (B) Mutations introduced in wildtype (wt) C^0^ clone S5206G5-42wk-c6.2 and wt C^2^ clone S5206G5-42wk-c3 to generate variants with altered V1 Cys content. (C) Longitudinal autologous plasma neutralization of wt and V1 Cys variant Envs. Heat map depicts the reciprocal half-maximal neutralization plasma dilution (NT50). Significance of neutralization differences was assessed by linear regression (ns = not significant). (D) Neutralization activity of heterologous bNAb-inducer plasma (7 donors, longitudinal timepoints) against S5206G5 V1 Cys variants shown in C. Heat map depicts the percent neutralization at 1/100 plasma dilution. Differences in neutralization sensitivities were assessed with linear mixed effects models to account for multiple plasma form the same donor in the analysis (ns = not significant, *** = p-value < 0.001). (E) Comparison of neutralization IC50s of bNAbs between C^0^ and C^2^ Envs derived from CATNAP. P-values are derived from Wilcoxon test by comparing median IC50 of each bNAb against the C^0^ and C^2^ Envs (** = p-value < 0.01, *** = p-value < 0.001).

To investigate the effect of the C^2^ motif on sensitivity to S5206G5 autologous plasma neutralization, we selected two e*nv* clones from week 42 that represented naturally occurring C^0^ and C^2^ genotypes and generated mutated versions by either removing or adding extra cysteine residues (Figure 3B). All Env variants had comparable autologous plasma NT50s (Fixed effects model p>0.05, Figure 3C). Both wt and mutant Env variants proved relatively resistant against heterologous bNAb-inducer plasma with different bNAb specificities (Table S6) with only few of the longitudinal plasma-virus combinations reaching NT50 >1:100 (Table S7). Comparing percent neutralization at a 1:100 plasma dilution across all longitudinal heterologous plasma profiles, we saw a significantly decreased plasma neutralization of C^2^ variants, compared to C^1^/C^0^ variants (Figure 3D) (linear mixed effects model p<0.001). The differences in plasma neutralization susceptibility were subtle, and most prominent in plasmas with weak neutralization activity such as S30834, S51928, S50725, S5213G5 and S10437 against Env S5206G5-42wk-c3.

We also measured the sensitivity of the five variants to a panel of reference bNAbs (N=14) (Table S3). The IC50s of bNAbs were not affected by the C^2^ motif (Wilcoxon p > 0.05) (Figure S2D) but similar to plasma inhibition, modest resistance increases between C^0^ and C^2^ inhibition were evident when comparing neutralization at a fixed, low bNAb concentration (Figure S2E). Taken together, these results indicate that while V1 Cys in these bNAb-inducer Envs have no clear impact on potent, high-dose bNAb-mediated neutralization, the C^2^ motif reduces neutralizing sensitivity under conditions such as lower bNAb doses or in the presence of weaker neutralizing plasma and antibodies.

To analyze the effect of the C^2^ motif on bNAb neutralization in diverse Env backgrounds, we harnessed neutralization and sequence data available on CATNAP^25^ for 21 bNAbs and 1260 *env* sequences that were either C^0^ (N=1189) or C^2^ (N=71). C^2^ Envs were significantly more resistant against V2-Apex bNAbs PG16, PG9, PGT145, V3-Glycan bNAb PGT121, CD4bs bNAb VRC01 and interface bNAb 35O22 (Wilcoxon p-values<0.01) (Figure 3E). The higher neutralization of C^2^ Envs in the CATNAP database suggests C^2^ insertions play a role in resistance development to bNAbs targeting diverse epitopes, in diverse Env backgrounds.

### Twin V1 Cys motif shifts accessibility of bNAb epitopes

We next hypothesized that C^2^ motifs may facilitate V1-mediated shielding by stabilizing the V1. To test this, we generated soluble C^2^ and C^1^ native-like SOSIP and DS-SOSIP trimers using a combination of previously reported stabilizing mutations (Table S8). We chose Env S5206G5-42wk-c3 which naturally carries a C^2^ motif and made two C^1^ versions of it by mutating either of the two extra V1 Cys to Ser (C146eS or C137S, HXB2 numbering). We then compared the binding profiles of the wild-type C^2^ and the two C^1^ variant trimers in a binding antibody multiplex assay (BAMA) against a panel of reference bNAbs and nAbs (N=29) (Figure 4, Figure S3, Table S3). Comparing the raw BAMA signals (mean fluorescence signals, MFI), removal of either Cys lead to enhanced binding of many bNAbs to C^1^ DS-SOSIP (Figure 4A-B) and C^1^ SOSIP (Figure S3A-B) trimers compared to the C^2^ versions. This was particularly marked for CD4bs, interface/fusion peptide, and several V3-glycan bNAbs whereas most V2 apex and selected V3-glycan bNAbs bound better to C^2^ Envs. mAbs 17b and 48d failed to bind DS-SOSIP trimers but showed equal capacity to bind SOSIP C^1^ and C^2^ variants (Figure S3A-F) confirming that the changes in epitope exposure upon removal of extra V1 Cys did not result in general trimer opening.

**Figure 4:**
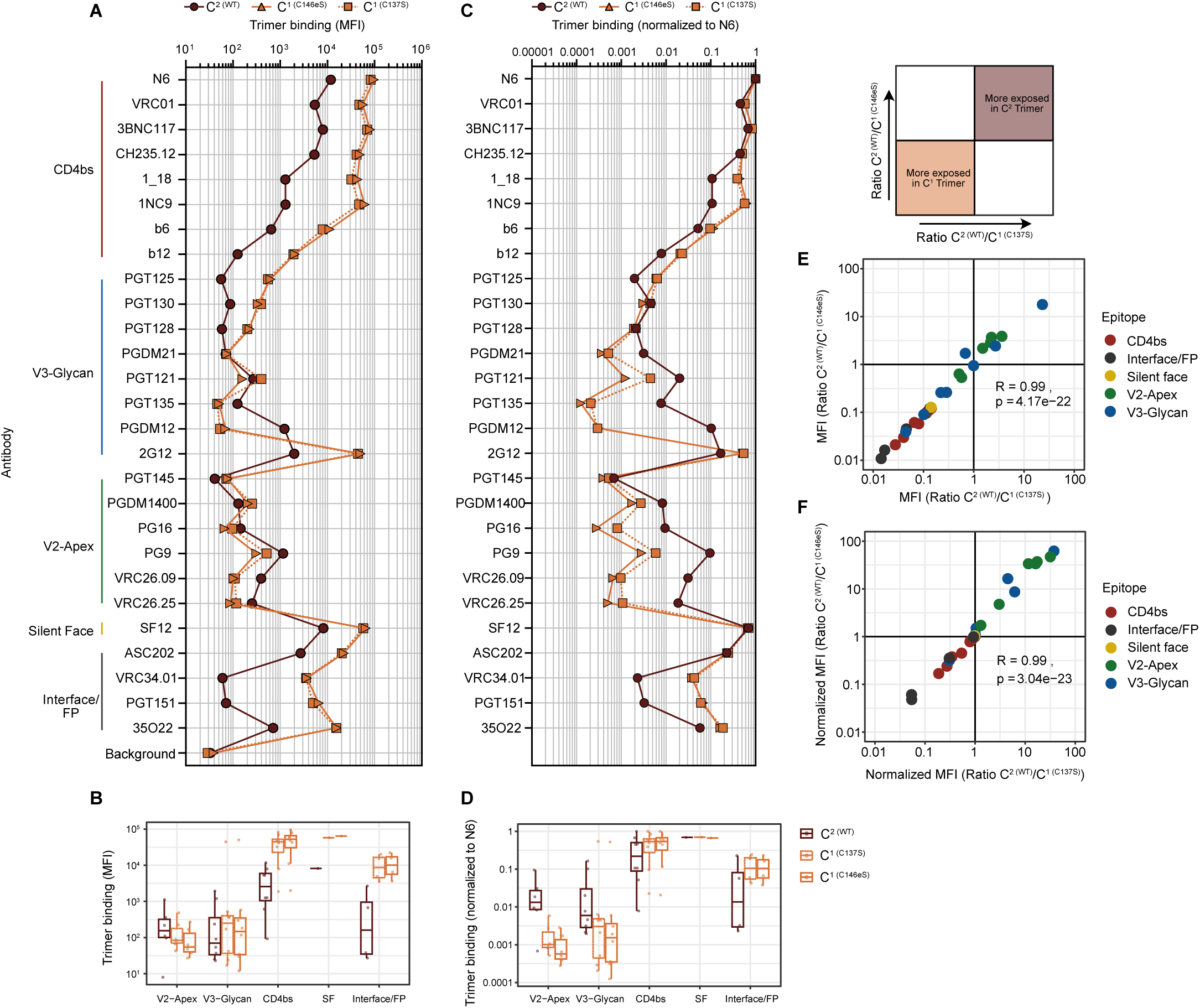
Extra V1 Cys modulate bNAb epitope exposure. (A) Luminex bead-based binding antibody multiplex assay (BAMA) of bNAbs (N=27) against soluble DS-SOSIP trimers of S5206G5-42wk-c3 Env variants: WT trimer (C^2^), with a C137S mutation (C^1^) and with a C146eS mutation (C^1^). Data depict mean fluorescence intensity (MFI). (B) Boxplots depicting the MFI values from (A) with antibodies grouped by epitope. (C) MFI values shown in (A) normalized to bNAb N6 which achieved the highest MFI signal in all data sets. (D) Boxplots depicting N6-normalized MFI values with antibodies grouped by epitope. (E) (F) Proportional differences in binding of C^2^ compared to C^1^ trimer versions based on data shown in (A) and (B) respectively. Each dot represents one bNAb, colors denote the bNAb epitope. Pearson correlation results are shown.

In order to define the relative accessibility of the bNAb epitopes in a given Env to each other, we normalized the BAMA data to the maximum binding observed for N6 (Figure 4C-D, Figure S3C-D). These analyses highlighted an increase in V2-Apex bnAb binding to C^2^ compared to C^1^ variants both by comparing effects on binding of individual bnAbs and by comparing on distinct regions (Figure 4D, Figure S3D). The alterations in epitope exposure from C^2^ to C^1^ were similar irrespective of which site was removed or whether we analyzed raw or normalized MFI (Figures 4E-F, S3E-F), suggesting that both Cys residues contribute equally to the epitope-modulating properties of the C^2^ motif, most likely by forming a disulfide bond.

Taken together, these findings indicate that the V1 Cys insertions in S5206G5 may have resulted from an escape from early autologous nAb activity suggesting that by inadvertently exposing the V2, the C^2^ motif may have generated an improved Env antigen that favored V2 bNAb evolution that occurred later in this donor.

### V1 Cys insertions are enriched among Env of bNAb-inducers

The high frequency of Envs with extra V1 Cys in the bNAb-inducer Env panel, along with the functional data, prompted us to investigate the role of the extra V1 Cys in the larger SHCS and ZPHI cohorts, which included participants of the Swiss 4.5K screen. We determined the frequency of V1 Cys insertions among participants using NGS-derived consensus V1 *env* sequences from plasma viruses (N=6424 sequences from 4373 unique participants) generated in previous studies^32,33^ and V1 sequences derived from single *env* clones (N=750 from 77 unique participants) (Figure S4A, Table S9). V1 sequences were used to study the frequency of Cys insertions in the context of bNAb evolution (Figure 5A-B), across infection stages including early transmission (Figure 5C, Figure S4B) and population-wide time trends (Figure S4C).

**Figure 5:**
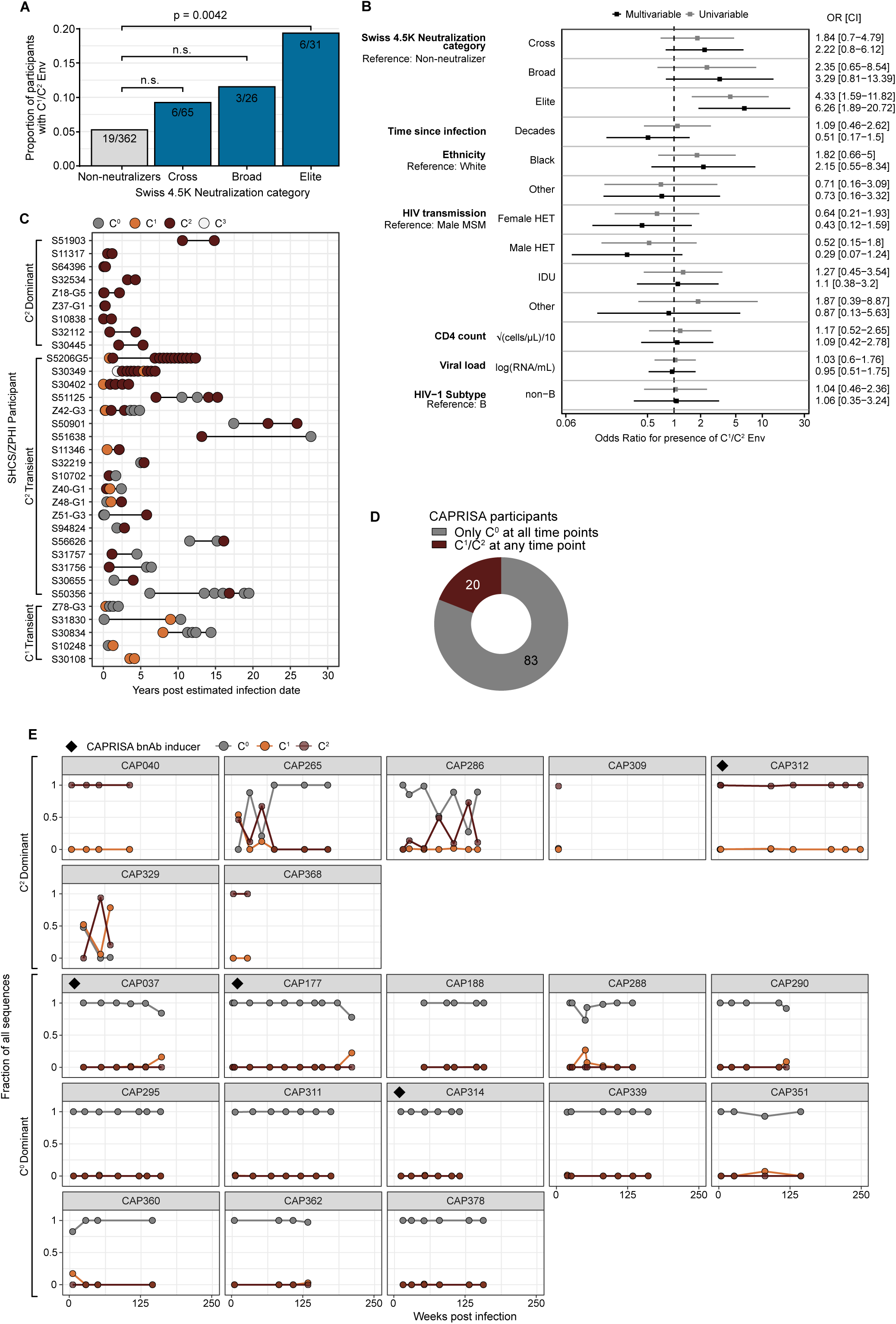
Non-canonical V1 Cys are enriched in bNAb-inducers. (A) Frequencies of donors with C^1^/C^2^ Env among 484 Swiss 4.5K participants with known neutralization status^19^. P-values calculated with logistic regression. (B) Uni- and multivariable logistic regression models assessing the independent contribution of host factors on the presence of V1 Cys insertions in Swiss 4.5K donors analyzed in (A). (C) Longitudinal V1 sequence analysis of 33 SHCS and ZPHI participants accounting for presence of C^0^, C^1^, C^2^ and C^3^ *env* in NGS consensus sequences or cloned *envs*. Each dot represents a time point. The dot color indicates the number of extra V1 Cys observed in the sequence. (D) Longitudinal V1 sequence analysis of CAPRISA donors (n=103) indicating the proportion of donor with (C^1^ or C^2^) or without extra V1 Cys detected at any time point based on single genome deep sequencing data. (E) Longitudinal frequencies of Envs with different V1Cys content for 20 CAPRISA participants with at least one C^1^ or C^2^ sequence. The proportion of observed V1 Cys variants is depicted.

For 484 of the 4,484 Swiss 4.5K participants with the required sequence data available (Figure S4A), we quantified the proportion of donors carrying *env* variants with extra V1 Cys residues as a function of their bNAb-inducer status (non-neutralizer, cross, broad and elite neutralizer) as defined in the Swiss 4.5K screen^19,20^. We observed a significantly higher frequency of V1 Cys in bNAb-inducers, with 6/31 elite neutralizers and only 19/362 non-neutralizers carrying viral variants with extra V1 Cys (19.4% vs 5.2%, Logistic regression, p = 0.0042) (Figure 5A). Using univariable and multivariable logistic regression models controlling for time since infection, ethnicity, transmission mode, CD4 count, viral load and HIV-1 subtype, we confirmed a positive, independent association between the presence of V1 Cys insertions and elite bNAb activity (adjusted odds ratio=6.26, 95% confidence interval (CI) 1.89,20.72) (Figure 5B).

We next probed the prevalence of extra V1 Cys at different disease stages by monitoring the frequency of V1 Cys insertions as a function of duration of untreated HIV-1 infection in 1528 SHCS and ZPHI donors (Figure S4A). We detected variants with V1 Cys insertions at all disease stages, including very early after transmission, were 7.5% of participants sequenced <90 days post infection had *env* variants with extra V1 Cys (Figure S4B). The frequency of participants with extra V1 Cys was constant throughout early and late infection ranging between 6.2-9.9% (Logistic regression p>0.05), suggesting that V1 Cys insertion is not restricted to specific disease stages (Figure S4B).

Since extra V1 Cys are associated with increased resistance to neutralization, we probed whether HIV-1 was evolving toward a higher frequency of extra V1 Cys at the population level. We restricted the analysis to sequences collected within the first year of infection to focus on viral variants that have not yet undergone extensive autologous neutralization escape. Analyzing SHCS and ZPHI Env sequences (N=431) sampled between 1996 to 2019 that met this criterion, we observed no significant differences in frequencies of participants with extra V1 Cys in between different sampling years (Logistic regression p>0.05) (Figure S4C). In a similar analysis, we examined the Los Alamos HIV Sequence Database^34^ for differences in year of sampling of C^0^, C^1^ and C^2^ sequences among viruses sampled <1 year post-seroconversion. There were no significant differences in sampling year, consistent with the SHCS and ZPHI cohort data (t-test p>0.05) (Figure S4D).

Longitudinal sequences were available for 33 SHCS and ZPHI participants identified as extra V1 Cys carriers allowing us to study within-donor temporal dynamics of V1 Cys insertions. Across longitudinal samples, C^0^ and C^2^ variants were the most common, with C^1^ and C^3^ (containing three extra V1 Cys) only appearing sporadically (Figure 5C). For 9/33 participants, C^2^ variants were detected at all time points (C^2^ dominant, Figure 5C). Nineteen participants had C^2^ variants appearing transiently and five participants had transient C^1^ sequences without any C^2^ variants detected. Of note, as these analyses were based on consensus sequences and single Env clones, we cannot exclude the possibility that V1 Cys variants persisted as minor variants for prolonged periods. A re-appearance of V1 Cys-containing variants is indicative of this scenario in participant S51125 (Figure 5C).

The consensus sequences and *env* clones of the SHCS/ZPHI provided information on whether or not extra V1 Cys were present at a given time point, but not their frequencies. To obtain further insights into the evolutionary dynamics and proportions of viral variants carrying extra V1 Cys, we analyzed longitudinal *env* V1 sequences from 103 women enrolled in the CAPRISA 002 cohort^35,36^. These sequences were obtained from deep sequencing of plasma virus sampled from very early infection (2 weeks) to 4 years post infection using the Pacific Biosciences SMRT (PacBio-SMRT) UMI approach^37^. This method adds a unique molecular identifier to each viral RNA molecule enabling a proportional analysis of C^0^, C^1^ and C^2^ *env* variants at each time point^38^. We analyzed an average of 191 sequences per participant at each time point (range 1-893) and found that 20 (19.4%) of the analyzed participants had at least one C^1^ or C^2^ sequence at any time point (Figure 5D). C^2^ Envs reached frequencies of >50% in seven donors (C^2^ dominant), one bNAb-inducer (CAP312) and six non-neutralizers (CAP040, CAP265, CAP286, CAP309, CAP329 and CAP368) (Figure 5E). In the remaining 13 participants C^0^ Envs dominated, with only transient increases in the frequencies of C^1^ Envs.

Taken together, the low frequency of C^1^ and C^2^ *envs* supports the notion that this motif may be associated with transient viral escape variants, and mostly confer an evolutionary advantage only during a short period.

### Over-representation of twin Cys among non-canonical V1 insertions

We next assessed whether selective forces drive insertion of extra Cys on a population level. Using *env* sequences from the Los Alamos HIV Sequence database (n=6657 sequences from unique PWH), we investigated the preference for one to four extra Cys insertions in V1. Among the 6657 sequences, 79 V1 regions were C^1^ (1.2%), and 356 V1 regions were C^2^ (5.3%). C^3^ and C^4^ sequences were rare (<0.12%) (Figure 6A). We observed a significant, albeit weak correlation between the number of Cys residues and the length of the V1 hypervariable region (Kendall’s tau = 0.167, p = 1.2×10^-60^), suggesting V1 length is only partially responsible for the insertion of Cys residues (Figure 6A). Comparing the fractions of C^0^, C^1^, C^2^, C^3^ and C^4^ sequences between different subtypes confirmed that, across all subtypes except F, C^0^ sequences were the most common followed by C^2^ (Figure 6B). The frequencies of sequences with extra V1 Cys were comparable between most subtypes except subtype G which showed an astonishingly high frequency of C^2^ sequences (35/116 sequences, 30.2%) (Figure 6B). We investigated whether the high frequency of C^2^ sequences in subtype G could be due to sampling biases. Out of the 116 analyzed subtype G sequences, sequences from Nigeria were much more likely to be C^2^ sequences than C^0^ sequences, as compared with sequences from other countries (Fisher’s exact test, p-value = 0.0032), suggesting that the high frequency of C^2^ Envs in subtype G may be due to limited sampling of geographically related sequences. Overall, these data illustrate that enrichment of the C^2^ motif is possible.

**Figure 6:**
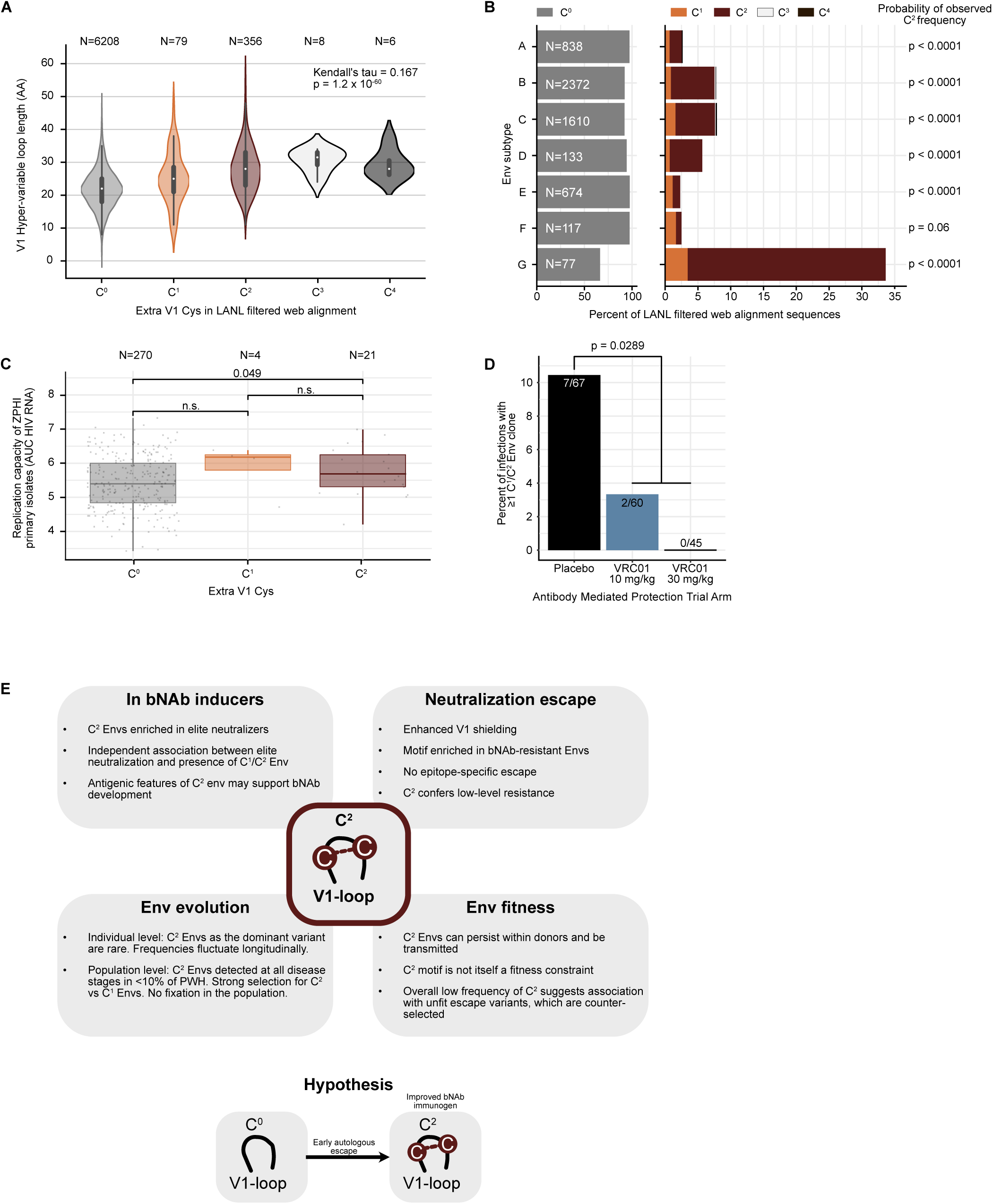
Fitness and selection of V1 with noncanonical Cys. (A) Correlation analysis of the length of the V1 hypervariable region and the number of Cys in V1 using the Los Alamos National Laboratory (LANL) HIV Sequence Database filtered web alignment (n=6657 *env* sequences) (B) Frequencies of C^0^, C^1^, C^2^, C^3^ and C^4^ Env sequences in different HIV-1 subtypes in the LANL HIV sequence database filtered web alignment. V1 hypervariable loop sequences were simulated for each subtype, 10000 times. P-values show the probability of observing at least as many C^2^ sequences in the simulated dataset as in the filtered web alignment. (C) Comparison of the replication capacity data of C^0^, C^1^ and C^2^ primary HIV isolates (N=257) sampled in acute to recent infection. Infectivity and sequence data are from the ZPHI cohort published in^39^ P-values were calculated using a student’s t-test. (D) Analysis of C^1^/C^2^ occurrence among 172 participants who acquired HIV-1 by the primary endpoint of 80 weeks in the HVTN703 and HVTN704 (AMP) trials probing bNAb VRC01 for prevention. Fraction of participants with at least one C^1^/C^2^ sequence in the placebo, low and high dose of VRC01 treatment arm is shown. P-value comparing Placebo to pooled VRC01 arms was calculated using Fisher’s exact test. (E) Summary of findings on the effect of V1 Cys insertions.

C^2^ sequences were strongly overrepresented compared to other categories of extra V1 Cys in all subtypes except F (Figure 6B). To analyze the probability of the observed C^2^ frequencies occurring by chance, we performed sequence simulations and generated distributions of C^1^, C^2^, C^3^ and C^4^ V1 sequences, assuming that Cys are randomly inserted with a probability calculated on their overall prevalence in the V1 of *env* (Figure S4E) (see Methods). The likelihood of observing at least as many sequences with two Cys in the simulated dataset was <0.0001, strongly suggesting that the presence of two Cys in the V1 hypervariable region is non-random and selected for (Figure S4E). This observation held true when limiting the analysis to sequences of a single subtype; except for subtype F, all subtypes showed a very low probability of similar C^2^ frequencies of twin Cys insertions in the V1 hypervariable loop occurring by chance (Figure 6B). The analyses of the LANL filtered web alignment suggest that there are selection pressures acting on the virus that select for the insertion of two Cys in the V1 hypervariable region.

### Comparable entry capacity and transmission potential of Env variants with extra V1 Cys

Occlusion of conserved Env regions by additional C^2^ bonds, including bNAb epitopes, may suggest that C^2^ variants have a selective advantage for nAb escape. In contrast, the low frequency and often transient nature of extra Cys variants indicates this motif may be associated with fitness deficits. To explore whether extra V1 Cys residues affect virus entry, we first studied the entry capacity of five Env clones of participant S5206G5 with different V1 Cys content naturally co-circulating at 42 week (Figure 3, Figure S2). When we examined the entry capacity of wild-type Envs (three C^2^, one C^1^ and one C^0^ clone) and matched mutant Envs, we did not find a consistent, strong effect of V1 Cys content on pseudovirus infection of TZM-bl cells (<4-fold changes between C^0^, C^1^ and C^2^ Envs) (Figure S5). Thus, in the context of the S5206G5 bNAb-inducer Env, the extra V1 Cys motif is well tolerated, consistent with its existence in later Envs (Figure 3).

We hypothesized that the lack of fitness costs in chronic Envs from a bNAb donor (who would have experienced unique selection pressures) may not reflect overall fitness deficits that are consistent with the low population level prevalence of this motif. We therefore used infection data from primary blood mononuclear cells (PBMC) of 295 primary, replication-competent plasma virus isolates from the ZPHI cohort, sampled in acute and recent infection^39^ when autologous neutralization is low. Of the 295 isolates which likely represent transmitted viruses, 7.1% were C^2^, 1.3% C^1^ and 91.5% C^0^ and all groups had comparable replication capacity (Figure 6C), confirming that viruses with extra Cys are fit and transmissible.

Lastly, we assessed whether putative fitness costs might be relevant only in the context of stronger selection immune selection pressure, such as that of bNAb prophylaxis. To address this, we used sequences from the Antibody Mediated Protection (AMP) trials HVTN703 and HVTN704^40,41^, in which the preventative capacity of the CD4bs bNAb VRC01 was assessed. In the two trials combined, 4611 participants were enrolled in three arms, 10 mg/kg VRC01 (N = 1537), 30 mg/kg VRC01 (N = 1539) and placebo (N = 1535). We assessed *env* clones from 172 participants who acquired HIV by the 80-week endpoint^40–42^. In total, 346 single genome Env clones (1-6 clones per participant) were derived from samples very close to infection (≤ 28 days post last negative test). Nine participants (5.2%) had at least one C^1^ or C^2^ *env* sequence (Figure 6D). Notably, seven of these participants were in the placebo arm, two in the low dose VRC01 arm and none in the high dose VRC01 arm suggesting a reduced transmission capacity of C^1^ and C^2^ Envs in the presence of VRC01 (Fisher’s exact test, p-value = 0.0289 for placebo versus combined VRC01 arms). Collectively, these data confirm that C^1^ and C^2^ Envs are transmissible to a new host where no neutralization pressure exists, but do not persist in the presence of strong selective pressure, accounting for their low overall prevalence (Figure 6E).

## Discussion

Defining Env genetic features enriched in bNAb-inducers is of interest for bNAb vaccine development as such features may contribute to the co-evolutionary process of bNAb elicitation. In previous research, we demonstrated that *env* genetics influence the specificity of the bNAb activity^19,20^, and that distinct Envs may harbor the capacity to elicit identical bNAb activity across individuals^24^. Translating this virus-encoded heritability of antibody responses to vaccine immunogens will be key to the success of HIV vaccines.

The identification of common HIV-1 Env signatures in bNAb-inducers is challenging due to the immense genetic diversity of *env*. In particular, the V1V2 region shows high genetic diversity while simultaneously being an important regulator of overall neutralization sensitivity^1,2,4^. Here, utilizing extensive data from the longitudinal SHCS and ZPHI cohorts, we show that the insertion of non-canonical Cys residues in V1 associates with elite bNAb induction. The insertion of Cys residues into V1 has previously been implicated as an escape strategy in some bNAb-inducers^6,29–31^. We extend this to show that additional V1 Cys are highly enriched in bNAb-inducers, and we confirm that this is a common strategy used by the virus regardless of bNAb status, infection stage and subtype.

We provide evidence for a multi-faceted role of extra V1 Cys in steering resistance to neutralization and in shaping bNAb activity (Figure 6E). HIV-1 selectively enriches for a twin Cys insertion in V1 (C^2^), as opposed to single Cys insertions (C^1^) on the population level, which we show to be linked with epitope shielding and modest increases in neutralization resistance. The over-representation of the C^2^ motif compared to C^1^ suggests that the formation of an additional disulfide bridge contributes to resistance. It is plausible that this extra bond provides stabilization to the V1, which may be particularly beneficial after extensive V1 elongation, as seen in certain bNAb-inducer Envs. Furthermore, as we show by analysis of the Los Alamos HIV Sequence database, the occurrence of V1 extra Cys is associated with an extended V1 length. However, the C^2^ motif also exists in Envs with shorter V1 loops suggesting that structural fixation of the V1 is not restricted to long V1 loops.

Selection pressures drive within-host evolution of HIV^43^. While escape from single bNAbs can be achieved by mutating few amino acids to inhibit bNAb-Env interactions, these mutations frequently come at a fitness cost for the virus^44,45^. In contrast to single escape mutations, our data suggest that resistance conferred by V1 elongation and C^2^ insertions is not directed to a specific epitope region; rather this appears to be a common strategy to increase resistance to multiple nAb epitope regions. The fact that the extra V1 Cys do not persist at high frequency in most donors suggests, however, that they are associated with other fitness deficits. We hypothesize that in the absence of an optimal escape mutation, the modest level of resistance conferred by the C^2^ motif may allow sufficient residual replication to acquire mutations that improve resistance and fitness.

We explored whether V1 Cys insertions negatively affect the entry process, leading to a reduced replication capacity of these viral variants. With increasing Cys content, the potential for aberrant disulfide bridging increases, which may affect the yields of correctly folded entry-competent trimers and therefore virus infectivity. Intriguingly, however, we did not observe strong entry capacity deficits linked to the C^2^ motif, and no significant differences in *in vitro* replication capacity of C^0^, C^1^ and C^2^ virus strains from a large number of transmitted founder (T/F) viruses from the ZPHI study^23,39^ were observed, suggesting that viruses with C2 motif can be transmitted. T/F Envs from a transmission pair carrying the C^2^ motif, which we previously documented (pair T6-R6 in^46^), are consistent with this observation. Data shown here from the CAPRISA cohort confirm that C^2^ Envs can be present very early in infection, underlining their transmission potential. Taken together, these results support the notion that the C^2^ motif is not itself a fitness constraint, but given its overall low prevalence, the motif may associate with unfit escape variants that are rapidly counter-selected.

Our longitudinal cohort studies investigated the role of the C^2^ motif in escape in an autologous immune environment relevant for evolution and transmission. However, in the context of bNAb vaccines or bNAb prophylaxis, the transmitted virus would face a new type of antibody, likely different to that from which it escaped. Indeed, the analysis of breakthrough cases in the AMP trials revealed that Envs with V1 Cys insertions appear to have a deficit in transmission when facing potent bNAb activity. This finding is consistent with the notion that additional V1 Cys confer a sufficient advantage to the virus only in a particular environment and at a particular time to become predominant. If these variants experience an altered immune environment in which their associated resistance patterns are no longer advantageous, e.g. due to newly emerging autologous nAbs or through bNAbs in prevention, as in the case of the AMP trial, fitness deficits will prevail, and they will be counter-selected (Figure 6E). However, we also present several examples of donors in which extra V1 Cys are maintained longitudinally including from early time points post infection onwards.

Viruses from bNAb-inducers that have co-evolved with an ever broader and more potent neutralization response are likely optimized for both fitness and resistance, increasing the likelihood that the added resistance provided by the C^2^ motif will remain important, be conserved, and become predominant. In support of this, two bNAb-inducers we previously defined as a transmission pair^24^, S30402 and S30349 (Figure 1A, Table S2), both had viruses with V1 Cys insertions present already in the early V3-glycan escape variant Env and both subsequently mounted a CD4bs bNAb activity^24^.

We hypothesize that the C^2^ motif may contribute to the bNAb imprinting capacity of Env by improving stabilization of trimer regions. We provide evidence for this scenario by studying the Env of the V2-Apex bNAb-inducer S5206-G5, which evolved to create a C^2^ variant, reducing access to several bNAb epitopes as expected, but increasing access to the V2-Apex region. This suggests that escape from early autologous neutralization activity necessitated a V1 elongation that was stabilized by the C^2^ motif. This introduction of C^2^ occurred at a stage when V2-bNAb activity had not yet evolved suggesting that the antigenic features of the C^2^ variant Env contributed to the evolution of V2-Apex bNAb activity in this donor. The role of autologous responses leading to V1 elongation and Cys insertion is supported by the fact that C^2^ *envs* were also detected in donors without cross-neutralizing activity.

Our data provide insights to the link between bNAb induction and insertion of V1 Cys. We describe two possible roles for the V1 Cys insertions in the context of bNAb development: they may aid autologous nAb/bNAb escape by shielding epitope regions and they may promote bNAb development by presenting a more stabilized immunogen. Both scenarios have implications for vaccine development. BNAb development in natural infection typically takes years and multiple rounds of co-evolution between B-cell receptor repertoires and Env are necessary. The association between the presence of Cys insertions in the V1 loop and bNAb induction may highlight an important intermediate step in the co-evolution, where pre-fusion Env is sufficiently stabilized for further maturation of bNAb lineages toward breadth. This, in turn, could be exploited in the design of boost immunogens to advance initial binding antibody responses toward neutralization breadth.

## Materials and Methods

### Human specimens

The study utilized biobanked plasma specimen of PWH identified as bNAb-inducers in the Swiss 4.5K screen^19,20^ for HIV-1 Env cloning and neutralization analyses. The specimens were obtained from the biobanks of the Swiss HIV Cohort study (SHCS)^21,22^ and the Zurich Primary HIV Infection Study (ZPHI)^23^. The SHCS is registered under the Swiss National Science longitudinal platform (http://www.snf.ch/en/funding/programmes/longitudinal-studies/Pages/default.aspx#Currently%20supported%20longitudinal%20studies). The ZPHI is an ongoing, observational, non-randomized, single center cohort founded in 2002 that specifically enrolls patients with documented acute or recent primary HIV-1 infection (ClinicalTrials.gov, identifier NCT00537966). The SHCS and the ZPHI were approved by the ethics committees of the participating institutions (Kantonale Ethikkommission Bern, Ethikkommission des Kantons St. Gallen, Comité Departemental d’Éthique des Spécialités Médicales et de Médicine Communautaire et de Premier Recours, Kantonale Ethikkommission Zürich, Repubblica et Cantone Ticino–Comitato Ethico Cantonale, Commission Cantonale d’Éthique de la Recherche sur l’Être Humain, Ethikkommission beider Basel for the SHCS and Kantonale Ethikkommission Zürich for the ZPHI), and written informed consent was obtained from all participants.

The study utilized biobanked plasma specimen of PWH with known bNAb-inducer or non-neutralizer status enrolled in the CAPRISA cohorts for V1 generated using the PacBio-SMRT UMI sequencing protocol^37^. The CAPRISA 002 Acute Infection cohort was established in 2004, and has been following women from early/acute stage of infection. Env sequences were truncated to the V1 hypervariable region to assess the number of Cys residues. The CAPRISA 002 study protocol was reviewed and approved by the Biomedical Research Ethics Committees of the University of KwaZulu-Natal (E013/04), the University of Cape Town (025/2004) and the University of the Witwatersrand (MM040202).

### Cells

HEK 293T cells (American Type Culture Collection, USA) and TZM-bl cells (NIH AIDS Reagent Program, USA) were cultivated in DMEM, supplemented with 10% heat-inactivated FBS, 100 U ml^−1^ penicillin and 100 µg ml^−1^ streptomycin (all from Gibco, Thermo Fisher Scientific, USA) at 37 °C, 5% CO2 and 80% relative humidity. Expi293F suspension cells (Thermo Fisher Scientific, USA) for protein expression were maintained in serum-free Expi293F expression medium (Thermo Fisher Scientific, USA), according to the manufacturer’s instructions. Cells were regularly tested for mycoplasma contamination and tested negative. No cell line authentication was performed.

### HIV-1 full-length Env cloning

HIV-1 full-length Env cloning was performed from undiluted bulk cDNA generated from plasma virus RNA as described^24^ and cloned into the pcDNA3.1/V5-His (Invitrogen, Thermo Fisher Scientific, USA) vector using the InFusion protocol (Takara Bio, Japan) according to the manufacturer’s instructions. Four to eight clones of each sample were picked from transformed XL10 gold ultra-competent cells (Agilent Technologies, USA), incubated in 5 ml at 37°C overnight in LB (Thermo Fisher Scientific, USA) and the plasmids were recovered with DNA minipreps (Qiagen, Germany). Sequencing libraries of plasmids were prepared with NexteraXT (Illumina, USA), sequenced using the MiSeq Reagent Kit v2 (50 cycles) (Illumina, USA) and Env sequences retrieved by EnvSeq (https://github.com/medvir/EnvSeq). Env cloned from Swiss 4.5K participants were utilized to create the bNAb-inducer pseudo-virus panel (N=34; Table S2). Additionally, V1 sequences were derived from Env cloned from SHCS participants (N=750 V1 sequences from 77 unique participants) to study extra cysteine content in V1 (Table S9).

### Env mutagenesis

Mutagenesis of Env expressing plasmids for pseudovirus production and for soluble Env production was performed using the QuickChange II XL kit (Agilent Technologies, USA) according to the manufacturer’s instructions using 10 ng of plasmid template encoding the respective Env, 125 ng of forward and 125 ng of reverse mutagenesis primers per reaction. Successful mutagenesis was confirmed with Sanger Sequencing (Microsynth, Switzerland).

### HIV-1 Env pseudoviruses

Env-pseudotyped viruses were prepared by co-transfection of HEK 293T cells with plasmids encoding the respective *Env* genes and the luciferase reporter HIV vector pNLluc-AM as described^47^. Input of Env pseudoviruses for neutralization assays was adjusted to yield virus infectivity corresponding to 5,000–20,000 relative light units (RLU; measured on a Dynex MLX reader, Dynex Technologies, USA) obtained by infection of TZM-bl cells in 96-well microtiter plates in the absence of inhibitors. Details on Env-pseudotyped viruses generated with corresponding GenBank entry and subtype is provided in Tables S2 and S4.

The 34 Envs included in the bNAb-inducer panel (Table S2) were isolated on average 438 weeks (8.4 years) post the estimated date of infection (EDI), the average calendar year of isolation was 2005. The 41-virus panel comprised Envs from both acute and chronic stage viruses including Transmitted /Founder (T/F) Envs and Envs form the global neutralization reference panel^48–50^. The average calendar year of isolation was 2005. For 35 Envs included in the 41-virus panel estimates on the infection stage at isolation were available: acute or acute T/F (N=7), <3 months (N=8), <12 months (N=12), chronic (N=8).

The information on the 123 CATNAP-chronic Envs was derived from the CATNAP database^25^ where they were specified as been isolated in chronic infection without further details on years post EDI. The average year of isolation of the CATNAP chronic Env panel was 2000 (Table S5). Overall, per reference bNAb, IC50 values for 14-114 of the total 123 analyzed CATNAP- chronic Envs were available (see Table S5 for the full listing).

### Neutralization assay

The neutralization activity of plasma and mAbs was tested on TZM-bl cells using Env-pseudotyped viruses in a 384-well format using a PerkinElmer EnVision Multilabel Reader (PerkinElmer, Inc., USA) to record luciferase reporter activity as described^47^. 4-fold serial dilutions of plasma and Abs were tested, with starting concentrations of 1:100 and 25 µg/ml, respectively. Concentrations correspond to the final assay volume. The low antibody dose recorded in Figure S2D corresponds to dilution 6 of the 3-fold serial dilution of Abs. The plasma or antibody concentrations causing 50% reduction in viral infectivity (half-maximum inhibitory concentration, IC50) were calculated by fitting data to sigmoid dose–response curves (variable slope) using Prism (GraphPad Software, Inc., USA). If 50% inhibition was not achieved at the highest or lowest inhibitor concentration, a greater than or less than value was recorded.

### bNAb plasma neutralization specificity

Neutralization specificity of bNAb plasma was defined through neutralization fingerprints of the plasma analyzed in the Swiss 4.5K screen using the fingerprinting method described there^19^. Briefly, plasma neutralization activity against a 41-virus multi-clade panel (Table S4) was compared to neutralization activity of 43 reference bNAbs (Table S3). For each plasma, the dominant plasma bNAb specificity was predicted using the maximal-Spearman-based prediction (MSBP) as described in^19^. Plasma neutralization specificity based on the fingerprint results are listed in Tables S1 and S6.

### V1 sequence analysis from SHCS and ZPHI participants

The V1 sequence analysis of SHCS and ZPHI participants (Figures 5A-C, S4B-C) included consensus V1 sequences from published data^32,33^ and V1 sequences from individual full-length Env cloned from Swiss 4.5K and SHCS/ZPHI participants (N=750 sequences from 77 participants) in this study (Table S9). Consensus V1 sequences (N=6424) were derived from near full-length HIV-1 genome next generation sequence (NGS) data sets of plasma virus from the SHCS and ZPHI cohorts that were generated by long overlapping amplicon sequencing on Illumina MiSeq^32,33^ followed by assembly with an in-house sequence alignment tool (available at https://github.com/medvir/SmaltAlign). Alignment was done with an initial *de novo* assembly assisted alignment to the HIV-1 reference genome HXB2, followed by three alignments against the iteratively improved consensus sequence^51^. From the final sequence alignment, the majority consensus sequence was generated with a depth threshold of ≥20 for each position. The respective nucleotide regions were extracted from each sequence using the local version of NCBI BLAST^52^. The BLAST database consisted of the appropriate regions from a panel of 459 reference sequences obtained from the Los Alamos HIV sequence database (https://www.hiv.lanl.gov/). To obtain the final consensus sequences on amino acid level, we made codon alignments of all blasted gene sequences using MACSE v2, to account for frameshifts, for the amino acid translation^53^.

For all analyses, we excluded sequences from participants without EDI estimate available, and sequences lacking the conserved Cys residues that flank the V1 hypervariable region (HXB2 positions 131 and 157) to exclude non-functional Env sequences. Criteria for sequences inclusion/exclusion in the respective analyses are depicted in Figure S4A. Consensus and clone-derived V1 sequences were compiled alongside parameters on infection length, time, bNAb status and demographic characteristics as available from the Swiss 4.5K screen^19^ or the SHCS and ZPHI databases for the analyses in Figs 5A-C, Figure S4B-C (see Table S9 for full source data).

To analyze interdependencies between bNAb status and occurrence of extra V1 Cys, we used V1 sequence data from Swiss 4.5K participants. To ascertain a relationship between V1 features and neutralization status, the analysis was restricted to sequences sampled within one year of the plasma neutralization measurement in the Swiss 4.5K screen^19,20^ that defined the bNAb status and that were collected a minimum one year post the EDI (Figure S4A). The selection criteria were fulfilled for 507 of the 4,484 Swiss 4.5K participants.

To probe the prevalence of extra V1 Cys at different disease stages by monitoring the frequency of V1 Cys insertions as a function of duration of untreated HIV-1 infection plasma neutralization data was not required, allowing a total of V1 sequences of additional SHCS and ZPHI donors to be included (total N= 1481 sequences, one per donor; Figure S4A).

### V1 sequence analysis of CAPRISA 002 participants

The V1 sequence analysis of CAPRISA 002^35^ participants with known bNAb-inducer status^54–56^ (Table S9, Figs 5D and E) included V1 sequences derived by PacBio UMI Env sequencing using a method modified from^37^ to generate products with single PCR, instead of nested PCR. See Table S9 for source data.

### V1 sequence analysis of AMP trial breakthrough viruses

The V1 sequence analysis of AMP trial breakthrough viruses^40,42^ (Figure 6D) included single-genome V1 sequences derived by PacBio UMI Env sequencing Env sequencing from^42^ made publicly available at https://atlas.scharp.org/cpas/project/HVTN%20Public%20Data/HVTN%20704%20HPTN%20085%20and%20HVTN%20703%20HPTN%20081%20AMP/begin.view.

### V1 sequence analysis of viruses in the Los Alamos HIV Sequence Database

HIV sequences from the 2021 Los Alamos HIV Sequence Database M group with CRFs Env amino acid filtered web alignment (alignment ID: 121P2) were analyzed, which consists of 6657 sequences, with only one sequence per PWH and with very similar sequences having been removed. Additional quality control steps were performed when generating this alignment, including removal of the following sequences: those defined as ‘problematic’ by the Los Alamos HIV Sequence Database, those with >1 frameshifts, and those with significant insertions and deletions (as assessed by manual curation).

To determine whether sampling biases were contributing to the high frequency of C^2^ sequences in subtype G, infection country was obtained from the Los Alamos HIV Sequence Database. For countries with at least 5 C^0^ and/or C^2^ sequences (Cameroon, Nigeria, and Spain), we performed Fisher’s exact tests on subtype G sequences comparing number of C^0^ and C^2^ sequences in each of these countries versus all other sequences. Statistics were Bonferroni corrected.

To assess the probability of the observed frequency of C^2^ sequences, sequence simulations were performed. For each sequence in the filtered web alignment, an equally long V1 hypervariable region sequence was simulated, randomly assigning Cys at each position, based on the overall frequency of Cys in all V1 sequences (P(C) = 0.00565). Simulations were repeated 10 000 times to generate a theoretical distribution of sequences with extra Cys. P-values were calculated by comparing the frequency of C^2^ sequences in the simulated dataset with the observed frequencies.

### Soluble native-like Env trimer variants expression and purification

Codon-optimized sequences of strain S5206G5-42wk-c3 gp140 wild type comprising amino acids 31-664 (HXB2 numbering) and containing stabilizing mutations including the SOSIP or the DS-SOSIP mutations (Table S8) were custom synthesized (Thermo Fisher Scientific, USA) and cloned into the CMV/R expression vector in-frame with a C-terminal AviTag^57^. Env proteins were produced in Expi293F suspension cells (Thermo Fisher Scientific, USA) by transient transfection using a furin-expressing helper plasmid at a 3:1 ratio (w/w). HIV-1 Env proteins were purified from culture supernatants using *Galanthus nivalis* lectin agarose-resin (Vector Laboratories, USA) and C-terminally biotinylated using the BirA enzyme (Avidity Biosciences, USA) according to the manufacturer’s instructions. Subsequent size exclusion chromatography (SEC) (Cytiva, USA) on a Superdex 200 increase 10/300 GL was performed to derive the trimer protein fraction.

### Binding antibody multiplex assay

Binding of mAbs to soluble trimers was measured with a bead-based binding antibody multiplex assay (BAMA). MagPlex®-Avidin microspheres (Luminex) were washed before use in 100 µl phosphate buffered saline (PBS) with 2% FCS using a Magnetic Plate Separator (Luminex). Biotinylated AviTag HIV-1 envelope trimers (100 nM) were coupled to washed 5×10^5^ MagPlex®-Avidin microspheres in a total volume of 150 µl PBS/2% FCS. Coupled beads were stored at 4°C for up to 4 weeks before use in the assay. On the day of analysis, per bNAb tested, 1000 coupled beads were washed twice with 100 µl 2% FCS in PBS and resuspended in 100 µl 2% FCS in PBS. The different trimer-coupled microspheres were pooled and incubated with 5 µg/ml of bNAbs (IgG1) in a total volume of 100 µl at room temperature for 1h under continuous shaking at 750 rpm on a Heidolph Titramax shaker. After washing microspheres twice with 2% FCS in PBS, PE-labeled detector antibody (mouse anti-human IgG-Fc; Southern Biotech, Cat#9040–09, clone JDC-10; 1 μg/ml) was added and beads were incubated at room temperature for 1h on a shaker, followed by two wash steps with 2% FCS in PBS. Bound PE-detector Ab was recorded as mean fluorescence intensity (MFI) on a FlexMap 3D instrument (Luminex). A minimum of 50 beads per antigen and antibody were acquired to guarantee accurate MFI values.

### Entry capacity

The entry capacity of Env variants was assessed by measuring the infectivity of Env pseudoviruses on TZM-bl cells, normalized to the p24 content of pseudovirus stocks. Infectivity and p24 content were measured in parallel from the same transfection and measurements were performed in biological duplicates. For the infectivity measurements, pseudovirus serial dilutions were added to TZM-bl cells and luciferase activity (relative light units (RLU)) was measured three days post-infection using Luciferase assay reagents (Promega, USA). In parallel, pseudovirus serial dilutions were inactivated with 2% Empigen in PBS and p24 content of virus lysates was measure by ELISA as previously described^58^. Briefly, p24 was captured on anti-p24 antibody (D7320, Aalto Bioreagents, Ireland) coated plates and detected using alkaline-phosphatase-coupled antibody BC1071-AP (Aalto Bioreagents, Ireland). P24 content was then interpolated from a standard curve of recombinant p24 (AG 6054, Aalto Bioreagents, Ireland). The obtained infectivity (RLU/µL) values were normalized to the p24 concentrations (ng/µL) of all virus stocks.

### Replication capacity

Replication capacity of primary virus isolates form the ZPHI studied analyzed in Figure 6C were derived from Rindler et al^39^.

### Statistical analyses

Statistical analyses were done in R (version 4.3.1) and in GraphPad Prism (Version 10.1.0, GraphPad Software, Inc., USA). Linear mixed-effects models and linear regressions were fitted using the lme4 package in R^59^.

To compare two groups, we used logistic regressions adjusted on confounders when relevant (HIV-1 subtype, ethnicity, time since infection, viral load, CD4 levels and transmission mode). A Wald test with two-sided hypothesis was used to assess the significance of the estimated odds-ratio.

## Figure preparation

Figures were prepared either using R version 4.3.1 and the ggplot2^60^ package or in GraphPad Prism version 10.1.0 (GraphPad Software, USA). Figures were assembled and finalized in Affinity Designer (Serif Europe Ltd, United Kingdom).

## Declaration of generative AI and AI-assisted technologies in the writing process

During the preparation of this work, the authors used DeepL Translate, DeepL Write, ChatGPT 3.5 and Copilot for language editing and scite_ for literature search. After using these tools, the authors reviewed and edited the content as needed and take full responsibility for the content of the publication.

## Acknowledgements

This was work was funded by the following research grants to AT: Swiss National Science Foundation (SNSF) grant SNF 3147308_201266, Novartis Foundation for medical-biological Research 23B127. This study was co-financed within the framework of the Swiss HIV Cohort Study, supported by the SNF (#148522 and 201369 to HFG), by the small nested SHCS project 744 and 745 (to AT), the SHCS research foundation and by funds of the Yvonne Jacob Foundation to HFG. The funders had no role in study design, data collection and analysis, decision to publish, or preparation of the manuscript. The SHCS data are collected by the five Swiss University Hospitals, two Cantonal Hospitals, 15 affiliated hospitals and 36 private physicians (listed in http://www.shcs.ch/180-health-care-providers). PLM and CW are supported by the SA MRC Strategic Health Innovations Program, the National Institutes of Health and by the Bill & Melinda Gates Foundation’s Collaboration for AIDS Vaccine Discovery (CAVD, INV-036842). PLM is supported by the South African Research Chairs Initiative of the Department of Science and Innovation and the National Research Foundation (Grant No 98341). Chloé Pasin was supported by funding from the University of Zurich.

We thank Jacqueline Weber, Silvan Grosse-Holz and Cyrille Niklaus for technical assistance. We particularly thank the participants and the clinical and administrative staff of the ZPHI, SHCS, and CAPRISA cohorts for their decades of dedication, without which this research would not have been possible.

## Author contributions

MCH and AT conceived and designed the study.

MCH, PR, NF and AT designed experiments

MCH, PR, SK, MS, MMK, and DS Conducted experiments

MCH, MZ, PR, CP, JM, MMK, NF, HM, KW, PLM, CW, RK and AT analyzed data.

KJM, AR, HK, CA, RT, MH, SY, KL, MP, RK, GD, SF, DHW, WD, ACD, MJ, SE, NM, NG, JIM,

CW, PLM, HFG, RDK managed patient cohorts, contributed patient samples and Env sequences and analyzed patient-related data.

MCH and AT wrote the paper, which all co-authors commented on.

## Declaration of interests

All authors declare no direct competing interest related to this study including financial interests, patents, board affiliations and paid consultant activities.

## Declaration of unrelated interests

AT has received unrelated unrestricted research grants from the SNSF, Bill and Melinda Gates Foundation, Gilead Sciences, Novartis Biomedical Research foundation, University of Zurich (UZH) foundation, UZH Clinical Research Priority Program, the SHCS, Fondation Dormeur, and honoraria from Roche Diagnostics and the Institute for biomedical research Bellinzona for consultant and scientific board activity and directs the Swiss National Reference Center for Retroviruses together with MH.

HFG has received unrelated unrestricted research grants from the SNSF, the SHCS, Yvonne Jacob Foundation, Bill and Melinda Gates Foundation, University of Zurich Clinical Research Priority Program, the National Institutes of Health, Gilead Sciences, ViiV healthcare and Roche. HFG has received personal fees from Merck, Gilead Sciences, ViiV, Janssen, GSK, Johnson and Johnson, and Novartis for consultancy or DSMB membership and a travel grant from Gilead. MH directs the Swiss National Reference Center for Retroviruses and has received unrelated unrestricted research grant from the UZH Clinical Research Priority Program, the SNSF, the SHCS, and the ETH PHRT. RDK has received unrelated grants from SNSF, the National Institutes of Health, and Gilead Sciences. KL has received unrelated research grants from the SNSF, the SHCS, the STCS, the ProPatient Foundation and the Bangerter Rhyner Foundation.

Within the last 5 years, KJM has received travel grants and advisory board honoraria from Gilead Sciences and ViiV; and the University of Zurich received research grants from Gilead Sciences and Novartis for studies that Dr Metzner serves as principal investigator.

The remaining authors declare no unrelated interests.

---

**Figure S1.**
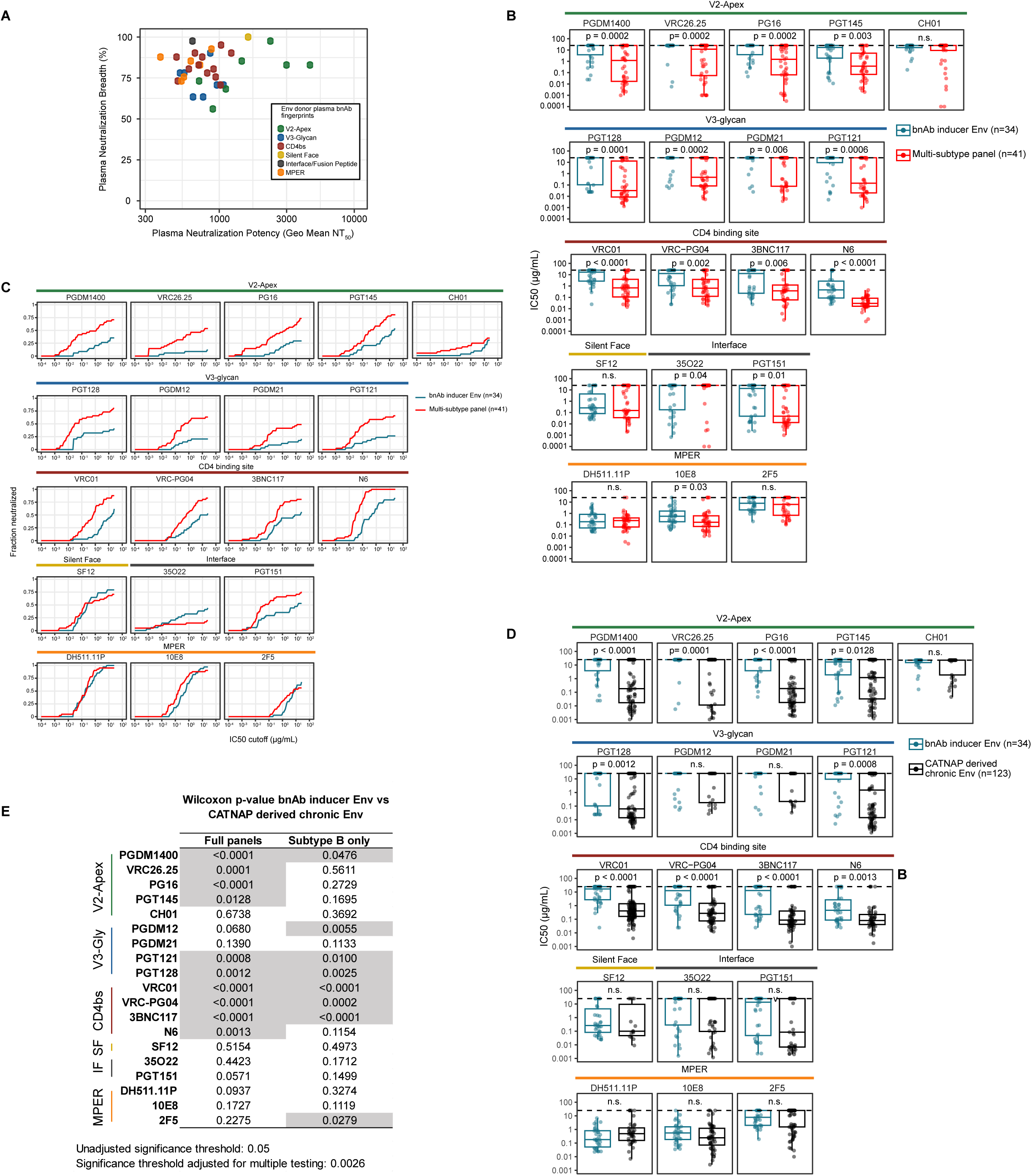
High resistance of bNAb-inducer Envs. (A) Plasma neutralization breadth and potency of bNAb-inducers whose Envs are included in the bNAb-inducer Env panel (n=34). Colors indicate the predicted plasma bNAb specificity. (B) IC50 of bNAbs against the bNAb-inducer Env panel (blue boxes) and the 41-virus multi-subtype Env panel (red boxes) (Table S2). Boxplots show the IC50 interquartile range, lines show the median IC50 and whiskers show the IC50 range. Points depict data of individual Envs. P-values indicate significance of the difference as assessed by Wilcoxon test. (C) Cumulative frequency distribution (breadth-potency curves) of the IC50 values of 19 bNAbs against the bNAb-inducer Env panel (blue) and 41-virus multi-subtype Env panel (red). (D) Comparison of bNAb IC50 against the bNAb-inducer Env panel (blue boxes) determined in the present study and bNAb IC50 reported in the CATNAP database for Envs isolated from chronic infection (CATNAP derived chronic Env). Boxplots show the IC50 interquartile range, lines show the median IC50 and whiskers show the IC50 range. Points depict data of individual Envs. P-values indicate significance of difference between the median IC50 of bNAbs in both Env panels as assessed by Wilcoxon test. (E) Wilcoxon test results depicted in (D) compared to the same analysis restricting to Subtype B Envs in in both Env panels.

**Figure S2.**
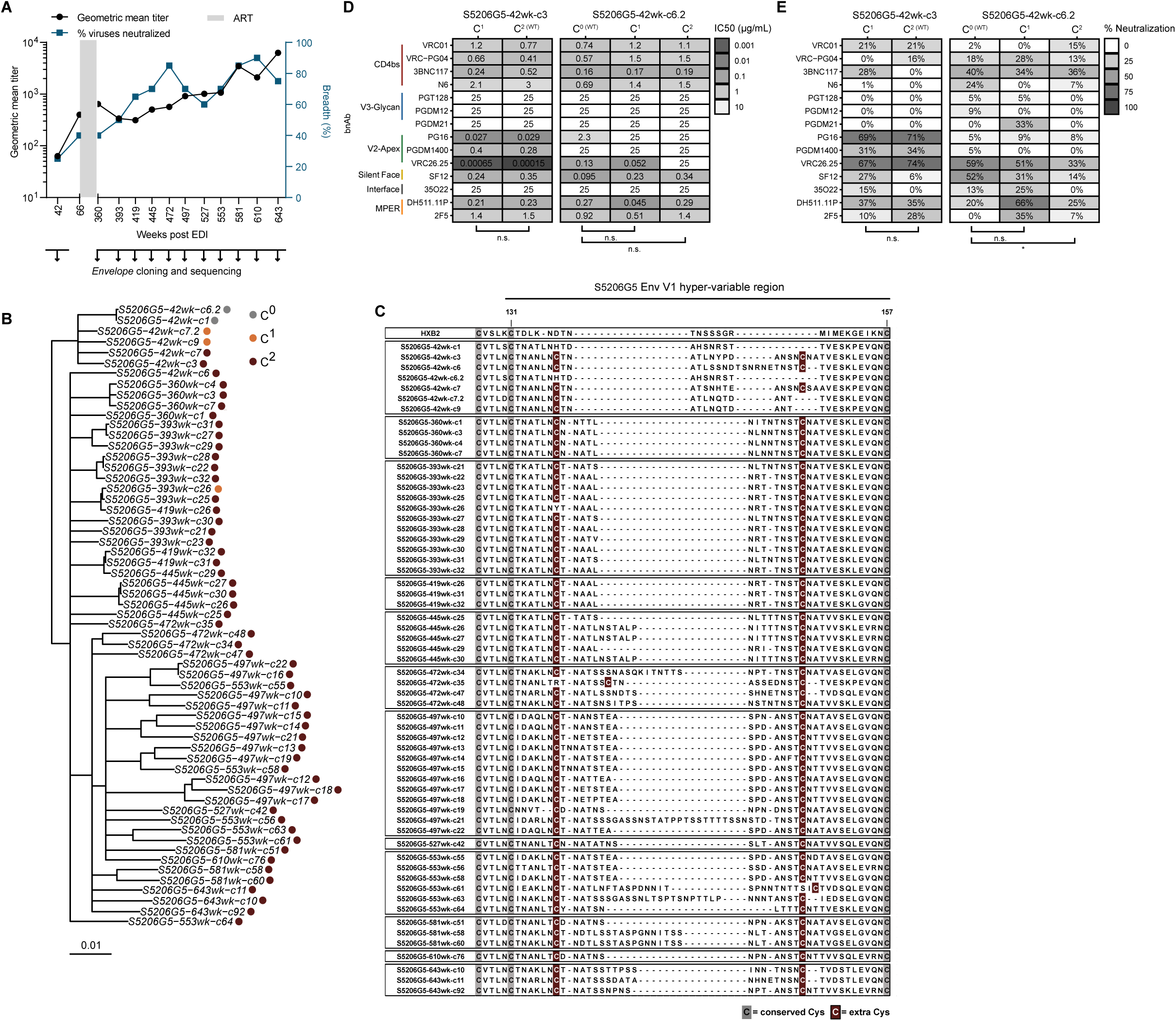
Modest impact of V1 Cys insertions on neutralization sensitivity. (A) Longitudinal plasma neutralization breadth and potency of S5206G5. Right: y-axis (blue) shows percentage of viruses neutralized from a 19-virus multi-clade panel. Left: y-axis (black) the geometric mean NT50 against the neutralized viruses. The gray area indicates antiretroviral therapy (ART). The arrows below the plot indicate time points from which Env was sequenced. (B) Phylogenetic tree of Env nucleotide sequences of the V2-Apex bNAb-inducer S5206G5 sampled 42-643 weeks post estimated infection date (EDI). C^0^ (gray), C^1^ (orange) and C^2^ (burgundy) circles indicate sequences with 0, 1 or 2 extra V1 Cys, respectively. (C) Multiple sequence alignment of longitudinal Env V1 hypervariable loop sequences from participant S5206G5. Conserved and extra V1 Cys residues are indicated by gray and burgundy shading, respectively. (D) Neutralization sensitivity of S5206G5 week 42 wildtype and V1Cys mutant clones. Heat map depicts IC50 of 14 bNAbs. Results of paired t-test are depicted (ns = not significant). (E) Data depicted in (C) displaying percent neutralization at a low bNAb concentration (0.103 µg/mL). Results of paired t-test are depicted (ns = not significant).

**Figure S3.**
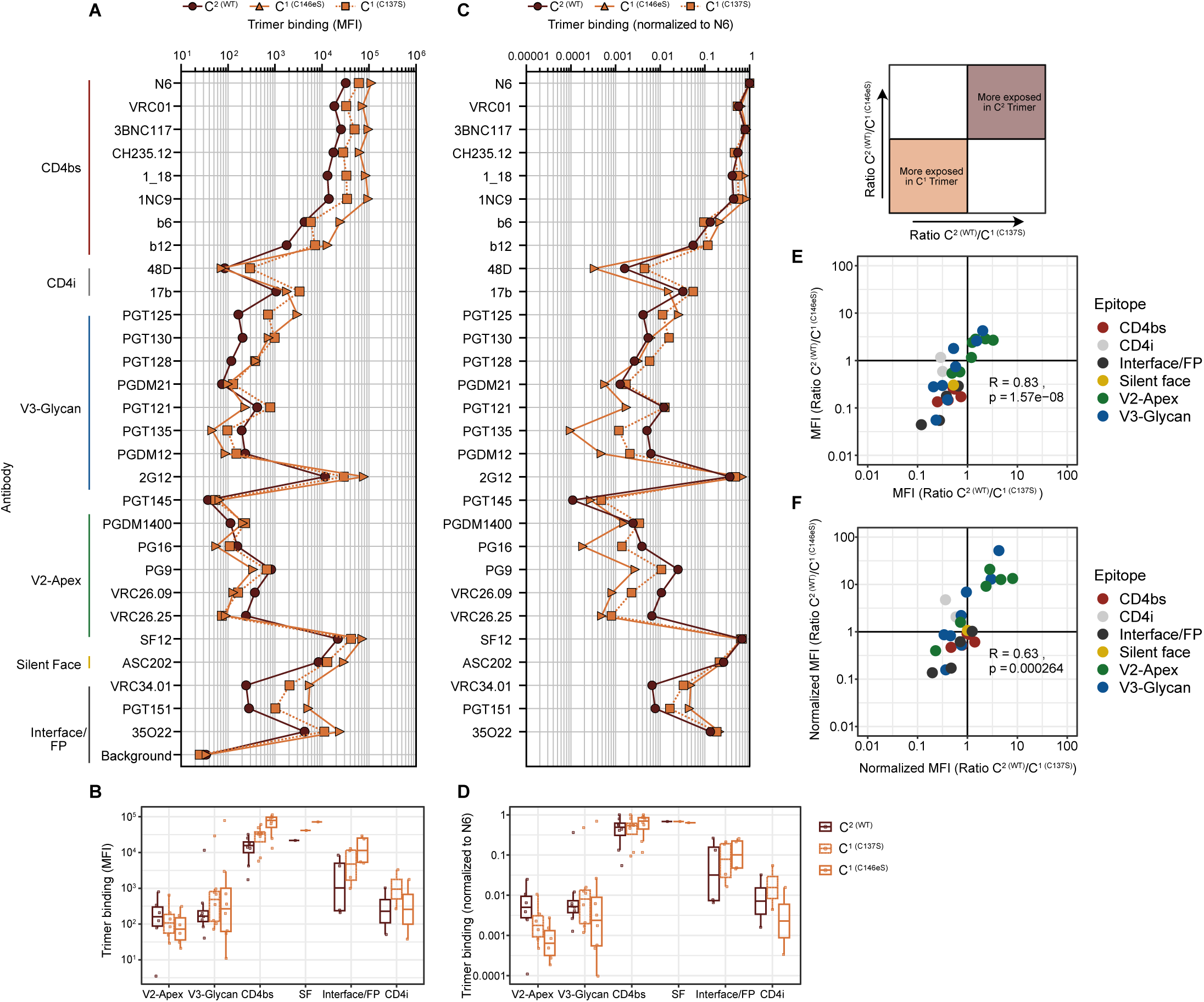
Antigenic properties of V1 Cys insertions. (A) Luminex bead-based binding antibody multiplex assay (BAMA) of bNAbs (N=29) against soluble SOSIP trimers of S5206G5-42wk-c3 Env variants: WT trimer (C^2^), with a C137S mutation (C^1^) and with a C146eS mutation (C^1^). Data depict mean fluorescence intensity (MFI). (B) Boxplots depicting the MFI values from (A) with antibodies grouped by epitope. (C) MFI values shown in (A) normalized to bNAb N6 which achieved the highest MFI signal in all data sets. (D) Boxplots depicting N6-normalized MFI values with antibodies grouped by epitope. (E) (F) Proportional differences in binding of C^2^ compared to C^1^ trimer versions based on data shown in (A) and (B) respectively. Each dot represents one bNAb, colors denote the bNAb epitope. Pearson correlation results are shown.

**Figure S4.**
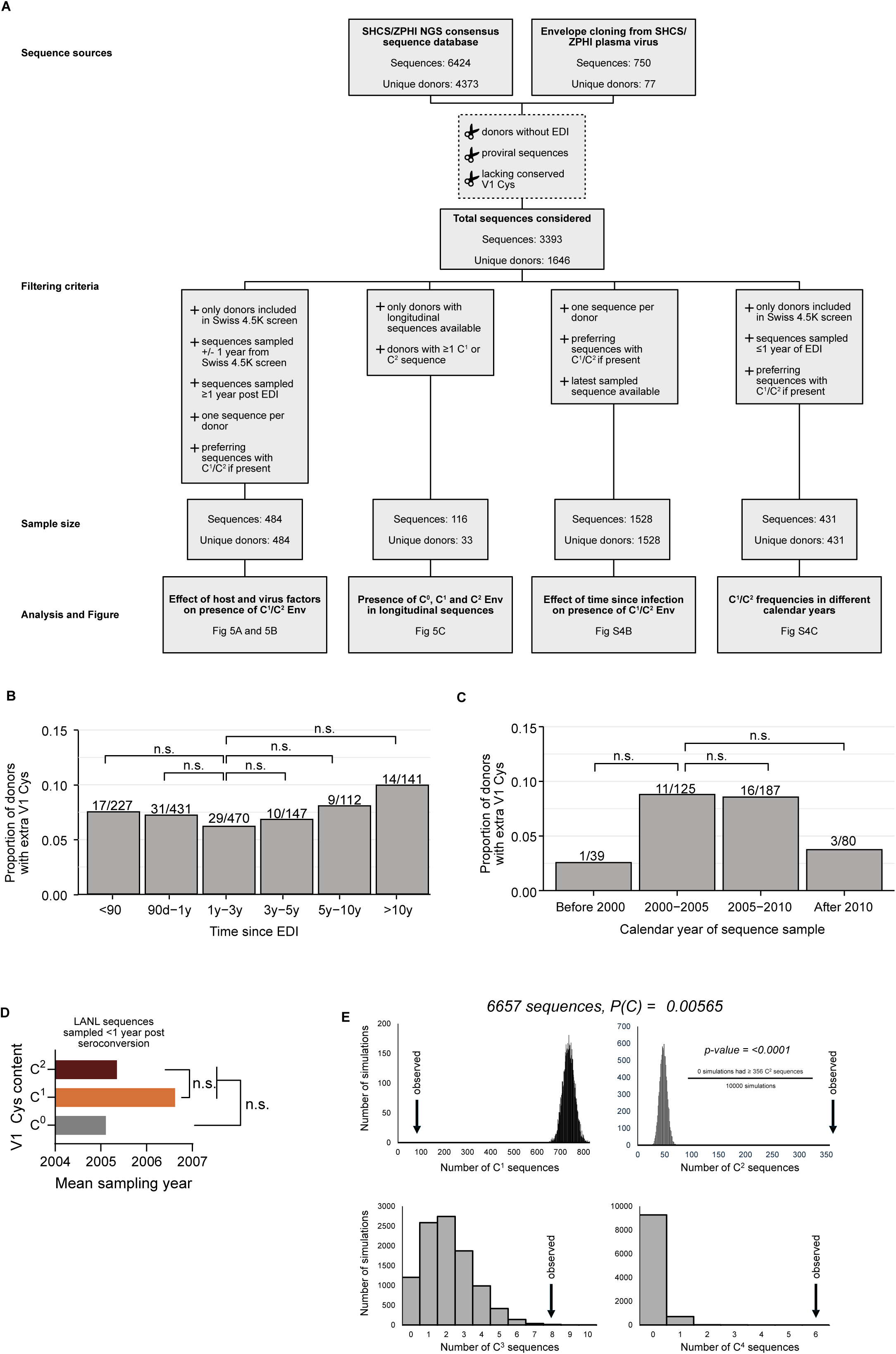
Researching extra V1 Cys at the population level. (A) Flow chart describing the selection of SHCS, ZPHI and Swiss 4.5K participants researched for V1 sequences in analyses depicted in Figs 5 A-C, S4B and C. (B) Proportion of 1528 SHCS/ZPHI participants with C^1^/C^2^ Envs depending on sampling time of the sequence in relation to the time since EDI. (C) Proportion of 431 SHCS/ZPHI participants with C^1^/C^2^ Envs depending on the calendar year of sequence sampling. (D) Mean sampling year of C^0^, C^1^ and C^2^ sequences in the LANL filtered web alignment. Differences assessed by t-test (n.s. = not significant). (E) For each sequence in the Los Alamos HIV Sequence Database filtered web alignment, an equal-length sequence was simulated. Whether or not a cysteine was present at each amino acid position was randomly assigned based on the overall prevalence of Cys in all V1 hypervariable sequences (P(C)=0.00565). Histograms depict the number of C^1^, C^2^, C^3^, and C^4^ sequences across simulations. The observed number of sequences of each type is labeled as “observed”.

**Figure S5.**
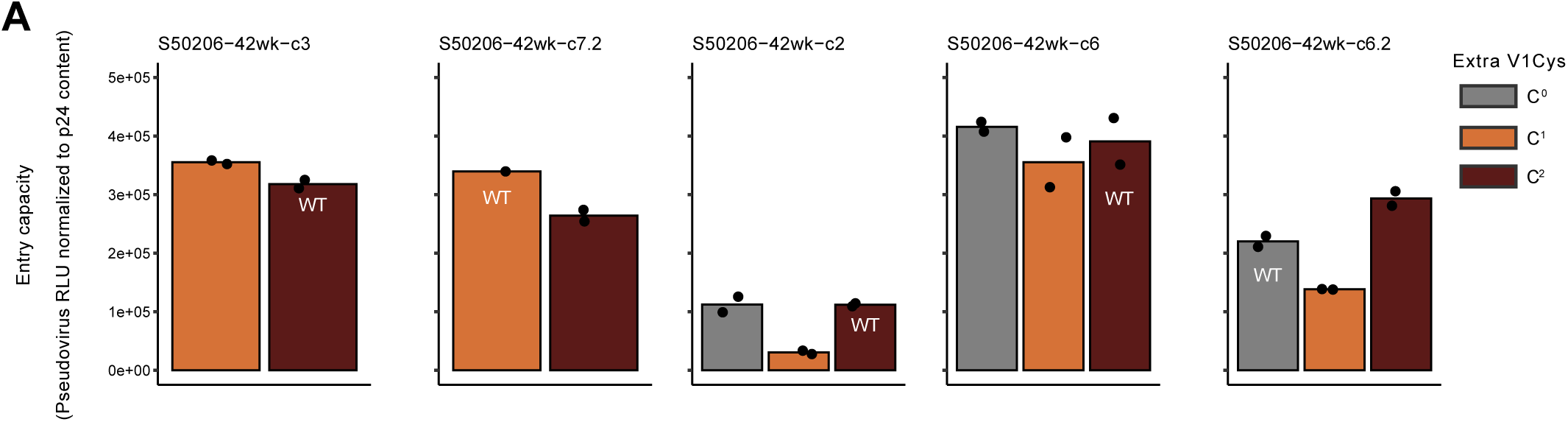
Entry capacity of V1 Cys variant Env. Entry capacity of S5206G5 Env V1 Cys variants was assessed by pseudovirus infection of TZM-bl cells normalized to p24 input. Relative light units (RLU) of emitted luciferase reporter activity, normalized to p24 content are shown as a measure of entry capacity.

## Notes

### Competing Interest Statement

The authors have declared no competing interest.

